# Progression of the cardiac phenotype of the DE50-MD dog model of Duchenne Muscular Dystrophy, corroborating results of cardiac magnetic resonance imaging with pathology up to 36 months of age

**DOI:** 10.1101/2025.02.28.640806

**Authors:** Julia Sargent, Virginia Luis Fuentes, Rebecca Terry, Dominique O. Riddell, Rachel CM. Harron, Dominic J. Wells, Richard J. Piercy

## Abstract

Cardiomyopathy is an expected consequence of the invariably fatal, X-linked muscle-wasting disease, Duchenne Muscular Dystrophy, (DMD) but onset and progression vary between individuals. Cardiac magnetic resonance imaging (CMR) is invaluable for identification of subclinical myocardial abnormalities, to stratify disease and to identify patients needing early therapeutic intervention. The dystrophin-deficient DE50-MD canine model harbours a mutation in the major hot spot region for mutations in DMD patients and mimics the preclinical cardiac and skeletal muscle of DMD boys. We performed serial parametric (T1, T2 and extracellular volume) mapping and late gadolinium enhancement (LGE) studies to characterise myocardial pathology in 15-to-36-month-old DE50-MD, (n=8) and wild type (WT, n=6) dogs. In 6/8 DE50-MD dogs left ventricular (LV) subepicardial LGE identified myocardial fibrofatty infiltration, (confirmed with histopathology post-mortem) with the characteristic distribution reported in DMD cardiomyopathy. Parametric mapping techniques exposed diffuse myocardial pathology that was most pronounced in DE50-MD dogs of 30 months and older and supported extracellular matrix expansion due to fibrosis, with or without regional myocardial oedema and /or fatty infiltration in individual dogs. All CMR markers of fibrosis worsened with age in DE50-MD dogs and paralleled deterioration in ventricular function evaluated with conventional and speckle tracking (circumferential strain) echocardiography, the burden of fragmented QRS complexes in 12-lead electrocardiography studies and the severity of pathological lesions. LGE and parametric mapping studies accurately identified a spectrum of myocardial abnormalities encountered in young DE50-MD dogs, further validating the relevance of these dogs as a preclinical model of DMD cardiomyopathy.

## Introduction

Duchenne Muscular dystrophy (DMD) is an invariably fatal, X-linked muscle-wasting disease. Absence of the protein dystrophin results in progressive skeletal muscle dysfunction, loss of ambulation by adolescence and ultimately, respiratory muscle failure. Concurrent myocardial injury in DMD patients leads to a distinctive pattern of progressive myocardial fibrosis, overt systolic heart failure, an end-stage dilated cardiomyopathy phenotype and poor prognosis.[1–9] Advances in supportive care and the use of palliative steroid treatment has increased longevity for DMD patients, and consequently cardiomyopathy has emerged as the leading cause of death.

The ability to detect and stratify myocardial damage is desirable to stage cardiac involvement, to identify individuals at increased risk of rapid disease progression and to guide decision making for therapeutic interventions. Characterisation of myocardial scarring using late gadolinium enhancement (LGE) imaging with cardiac magnetic resonance (CMR) has been extensively reported in DMD patients[2–5, 10–14] but has limitations: in particular, studies corroborating LGE imaging with specific histopathological findings in DMD patients are lacking. Further, the presence of LGE discriminates poorly between fibrosis, oedema and necrosis, all of which have been implicated in DMD-related cardiomyopathy.[15] Despite the recognised association between LGE and cardiomyopathy severity, global systolic dysfunction and regional wall motion abnormalities have been identified in patients without LGE. This could reflect the insensitivity of LGE to identify diffuse pathological changes, when gadolinium accumulation is lower and there is reduced contrast with the surrounding myocardium.[16, 17]

There is rising interest in the ability of cardiac parametric mapping techniques, (namely native T1, T2 and Extracellular volume (ECV) mapping) to provide quantitative characterisation of interstitial changes within the myocardium on a pixel-by-pixel basis. The T1 time, or spin-lattice time, is the time constant for the longitudinal recovery of magnetisation as it returns to an equilibrium state after the application of a magnetic pulse.[18, 19] It is therefore an intrinsic property of tissue that is altered by tissue composition and its interaction with neighbouring tissues. Myocardial T1 times are increased where there is increased interstitial water content; e.g. in the setting of myocardial inflammation/oedema and/or increased collagen content (i.e., fibrosis), as water is trapped within the expanded extracellular matrix (ECM). Myocardial T1 times are shortened, however, when there is fatty replacement of the myocardium, (e.g. in lipomatous metaplasia)[20] or following administration of gadolinium-based contrast agents.[21] Post-contrast T1 time is useful to evaluate fibrosis since gadolinium accumulates where there is extracellular matrix expansion, (resulting in regional shortening of T1 times)[22, 23] but is affected by the value of distribution and clearance of gadolinium, which can vary depending on individual patient factors and lead to inaccuracy.[24, 25] Calculation of ECV helps to ameliorate some of these confounding factors. Extracellular volume maps are generated following acquisition of native and post-contrast T1 maps and correction for red blood cell density according to the patient’s haematocrit. Extracellular volume fraction is a biological parameter that describes the non-myocyte proportion of the myocardium, encompassing the extracellular matrix and the myocardial blood volume.[19] It is consequently increased when there is expansion of the ECM; e.g. due to oedema or fibrosis, or infiltration by fat or amyloid [19, 26–28] and has been validated as a marker of myocardial fibrosis against results of histology in various cardiac diseases, though not in DMD.[29–32].

Several studies have identified that global T1 values are increased in DMD boys, supporting that diffuse ECM expansion is a pathological feature of DMD cardiomyopathy. Myocardial T1 values further stratify disease in DMD, such that patients with regional and focal ECM expansion, (i.e. those with LGE) and LV systolic dysfunction (i.e. those with a depressed EF) have the highest values.[33] The considerable heterogeneity of segmental T1 measurements within DMD individuals exceeds that of healthy controls, which likely reflects regional variation in the severity of myocardial injury.[34] The highest T1 values are present in the lateral wall segments of affected individuals and demonstrate more consistent differences in myocardial T1 values between LGE positive and negative patients,[33, 35] and those of LGE negative and age-matched, heathy boys.[33] Subepicardial T1 values exceed those of the subendocardial LV myocardium in DMD boys but not healthy controls, and show superior prediction of DMD severity and LGE status to global or subendocardial values.[34] The subepicardial layer of the mid and basal lateral wall segments is typically the earliest region of the myocardium in which LGE is identified in DMD cardiomyopathy, approximating the regional distribution of fibrofatty replacement described in DMD pathological cardiac specimens.[36, 37] These findings support that LGE represents a manifestation of severe myocardial injury along a diffuse spectrum of ECM expansion in DMD cardiomyopathy. Myocardial ECV is also elevated in DMD boys[35, 38, 39] but does not reliably stratify disease according to LGE status.[20, 40–42]

The T2 time is the time constant representing the decay of transverse magnetisation (spin-spin relaxation).[18] T2 mapping techniques allow regional quantification of the myocardial T2 on a pixel-by-pixel basis, using a similar principle to that used in T1 mapping, being elevated in the presence of fatty infiltration or oedema. Measured T2 times are elevated in the upper arm,[43] thigh and lower leg muscles of DMD boys[44–48] and the pelvic limb muscles of dystrophin-deficient canine models [49–51] due to the combination of oedema, inflammatory cell infiltration, and fatty infiltration of the skeletal muscle. The application of T2 mapping to dystrophin-deficient cardiomyopathy is less well described. Given that fibrofatty replacement is a recognised feature of DMD cardiomyopathy,[52] and recent concern regarding the possible impact of myocarditis in DMD boys,[13, 15, 53] results of T2 mapping could provide additional clues regarding myocardial characterisation in affected patients. To date, studies of a small number of DMD boys have demonstrated similar[43] or reduced[40, 54, 55] myocardial T2 times compared with results from healthy individuals. There can be marked variation between individual myocardial segments however, such that heterogenicity of T2 distribution is not only increased in DMD boys but is also predictive of a decline in LV ejection fraction (EF) and the myocardial deformation parameter, circumferential strain.[56]

Since the presence of LGE and elevation of T1 times does not accurately discriminate between fibrosis and oedema, T2 mapping can therefore provide additional information regarding myocardial composition in individual DMD patients akin to a non-invasive “virtual biopsy”. [38, 57] Validation of this strategy is challenging in the setting of DMD patients due to the lack of cardiac tissue available for histological corroboration. Translational animal models of DMD provide an invaluable opportunity to validate proposed biomarkers of disease progression against histopathological abnormalities; specifically imaging (transthoracic echocardiography (TTE) and CMR) and electrocardiographic features of the disease.

In this study we describe the use of parametric mapping techniques (T1, T2 and ECV mapping) to further characterise the myocardial composition of the dystrophin-deficient canine model of preclinical DMD cardiomyopathy, DE50-MD dogs. These dogs harbour a mutation that is located in the middle of the major hot spot region for mutations in DMD patients that results in deletion of exon 50 from DMD gene transcripts. The location of their mutation make them highly relevant to study novel treatments, particularly mutation-specific molecular treatments aimed at genetic repair[58] given that their mutation is relevant to the greatest number of human DMD patients.[59]Our previous work has demonstrated that, alongside their skeletal muscle and cognitive preclinical phenotype, [51, 60–62] these dogs exhibit an early preclinical cardiac phenotype that recapitulates that of dystrophin-deficient boys[63]. Herein, we explore the associations between the results of CMR imaging and those of conscious functional transthoracic echocardiographic (TTE) and electrocardiography (ECG) studies in young adult DE50-MD and age matched, male Wild Type (WT) dogs aged 15-36 months. Further, we compare results of CMR with postmortem histological assessment of DMD cardiac specimens.

## Materials and methods

### Study population

All dogs were recruited from the DE50-MD colony housed in a dedicated facility at the Royal Veterinary College (RVC), London as described elsewhere. [61, 62] Dogs in the study were produced after natural mating of carrier female Beagle (RCC strain, Marshall Bioresources)-cross (F3 generation) dogs derived from an original founder Bichon-Frise cross Cavalier King Charles Spaniel female carrier with healthy male stud Beagles (RCC strain). Dogs were cared for in conditions according to RVC local Animal Welfare Ethical Review Body approval and exceeding the minimum stipulations of the UK’s Animal (Scientific Procedure’s) Act, (A(SP)A) 1986. Dogs underwent an extensive socialization programme, with daily human interaction and acclimatization to routine procedures. They were given daily access to large outdoor paddocks (approximately 100 m2) in group sizes of up to 5 dogs. Adult dogs were housed in groups of 2 to 4 animals, according to temperament and hierarchy in large kennels (4.5m2-7.5m2, 12-hour light/dark cycle; 15–24 °C). Pregnant females were single housed when close to whelping, which was allowed to proceed naturally. Puppies were kept with their dams with heat lamps (∼28^0^C) until weaning and then grouped with their litter mates until 4-5 months of age. Puppies were weaned from 4 weeks of age, when they were allowed ad lib access to (Burns Original Chicken and Rice) puppy food and water. From 12 weeks of age they received 3-4 x daily feeds with Royal Canin Puppy ProTech Colostrum and Milk mixed with Burns Puppy Original Chicken and Rice and then transitioned to twice daily feeds of Burns Puppy Original Chicken and Rice mixed with Royal Canin Gastro-Intestinal Puppy Food (2:1) and ad lib water when over 6 months of age.

The study was designed following the ARRIVE guidelines and experimental procedures were performed within the stipulations of two successive project licences, (70/7515 and P9A1D1D6E) assigned under the UK’s A(SP)A 1986 and approved by the RVC’s Animal Welfare Ethical Review Board.

Puppies were microchipped at ∼ 1 week of age and subsequently genotyped from cheek swab DNA (GeneJET genomic DNA kit, Thermofisher) for creation and purification of PCR products (QIAquick, Qiagen) and Sanger sequencing (SupremeRun, Eurofins) as described previously[62] Disease status was corroborated though measurement of serum creatine kinase activity (data not shown). Male WT and DE50-MD puppies born into the colony were enrolled onto the natural history study from ∼15 months of age until the 36-month endpoint. Most dogs (5/8 DE50-MD and 6/6 WT dogs) were included in an earlier 3-to-18-month early cardiac phenotype study [63]. The three additional DE50-MD dogs being recruited at 15 months of age. Some carrier females were kept for colony maintenance (carrier females). All WT male dogs in this study, and animals not used for research, were rehomed. The latter included neutered carrier females and WT females and male dogs.

Strict attention was taken minimise any animal suffering throughout the study. Twice daily welfare assessments were conducted by animal technicians. Care was taken to identify any animals that could be approaching pre- determined endpoints for DE50-MD dogs; dehydration (unresolved by fluid treatment), lethargy/motor dysfunction, weight loss/dysphagia, dyspnoea, listless behaviour/demeanour, or heart failure. Any animal raising concern was reported to and assessed by the Study Director, the Named Veterinary Surgeon (NVS) and the Named Animal Care and Welfare Officer (NACWO). All DE50-MD dogs underwent humane euthanasia at the end of the planned 36-month study period or before, if they reached a pre-determined endpoint as detailed above (see results for additional detail). Euthanasia was performed following intravenous injection of sodium pentobarbital, (250 mg/kg, Dolethal, Covetrus).

### Blood Sampling

Blood samples were obtained from DE50-MD and WT dogs via jugular venipuncture at 6 monthly intervals between 18 and 36 months of age as part of a parallel study evaluating blood-borne musculoskeletal disease biomarkers in the DE50-MD model.[64] Approximately 2.5mls of blood was obtained in lithium heparin and plain collection tubes to obtain plasma and serum respectively. Blood samples were spun in a centrifuge at 500 x g for 10 minutes at 4°C. Serum was extracted from the blood samples, aliquoted into 1.5ml Eppendorf tubes and frozen at -80°C until required for analysis. Samples were sent to an external commercial laboratory (IDEXX Laboratories, Ludwigsburg, Germany) for quantification of cardiac troponin I (cTnI) using a high sensitivity assay (healthy range <70pg/ml) and N-Terminal pro brain natriuretic peptide (NT proBNP, healthy range <900pmol/l).

Values below the lower limit of detection of the NT-proBNP assay (<250pmol/l) were documented as 250pmol/l. Results of circulating biomarkers for dogs aged 18 months old were included in previously reported data [63].

### Antemortem cardiac evaluations

Serial cardiac assessments using TTE and 12-lead ECG assessments were scheduled with an intended study interval of 12 weeks. CMR imaging was performed at approximately 6-month intervals. Contemporaneous conscious TTE and electrocardiographic assessments were performed either within a week prior to, or between 48 hours and 7 days after recovery from general anaesthesia for CMR evaluation.

### Transthoracic echocardiography

Echocardiographic examinations were performed within 7 days of CMR examinations. Where TTE assessments were performed after CMR, examinations took place at least 48 hours after recovery from general anaesthesia. Conventional TTE echocardiographic and tissue Doppler imaging assessments were performed as previously described.[63]. All TTE examinations were performed and analysed by the same cardiology diplomate of the American College of Veterinary Internal Medicine using the same ultrasound machine (Vivid E9, General Electric Medical Systems Ultrasound, Hatfield, UK). All studies were analysed offline using proprietary software (Echopac, General Electric Medical Systems Ultrasound, Hatfield, UK.). Measurements were recorded as averages from at least 5 cardiac cycles. All standard images and measurements were collected according to the recommendations of the American Society of Echocardiography.[65] For a detailed description of measurements and echocardiographic protocols please see supplementary information. In brief, parameters of interest were weight-adjusted systolic and diastolic LV internal diameter (LVIDSN and LVIDDN respectively), ejection fraction (EF), Fractional shortening (FS) and Tricuspid Annular Plane Systolic Excursion (TAPSE). Tissue Doppler imaging of the basal LV free wall (FW), septum and right ventricle (RV) FW was performed to record longitudinal myocardial systolic velocities as described elsewhere[63]. Velocities were recorded from the left apical 4 chamber view using pulsed-wave spectral Doppler. Pulsed-wave sampling gates (2-3mm wide) were placed at basal annular segments to obtain the pulsed-wave spectral Doppler profiles.[66] Values for RS’ were adjusted for body size using the equation RS/BW^0.233^[67].

Speckle-tracking echocardiography derived circumferential strain parameters were derived from high-quality images collected by the same observer, using the same machine and transducer as for the conventional TTE assessment. Echocardiographic loops were stored from the right parasternal short axis view at the level of the LV papillary muscles to obtain circumferential deformation parameters.[68] Images were acquired, and image analysis performed offline to obtain peak systolic strain (S) and strain rate (SR) during systole (SSR), early diastole (ESR) and late diastole (ASR). Average peak strain and strain rate parameters were calculated by averaging the results from the 6 individual segments from 3 cycles. Circumferential deformation values were also calculated separately for the septum and LV FW separately by averaging results from the two midventricular septal and four FW segments, respectively.

### 12 lead electrocardiography

Twelve lead ECG assessments were performed immediately prior to echocardiographic assessment to obtain recordings from the frontal plane and chest leads [69, 70] (for detailed protocol please see supplementary information). Recordings for each lead were scrutinised for the presence of fragmented QRS complexes (fQRS). Fragmented QRS complexes were defined by the presence of one or more additional R waves (i.e. R’ waves) or notching in the nadir of the R-wave or the S-wave as described elsewhere.[71, 72] The presence of fQRS complexes had to be demonstrated in at least 2 contiguous leads to be considered present. The number of leads demonstrating fQRS was recorded for each dog at 3-month intervals.

### Cardiac magnetic resonance imaging

CMR imaging assessments (1·5 Tesla Magnetic resonance imaging (MRI) machine, (Intera Pulsar System, Philips Medical Systems, Surrey) and a flexible, sensitivity-encoding, phased array surface coil (SENSETM Flex M) were performed under general anaesthesia. All dogs received premedication with intravenous methadone (0.2mg/kg, Synthadon, Animalcare) prior to induction of anaesthesia with propofol, (4-6mg/kg, Propoflo, Zoetis) and maintenance with sevoflurane, (SevoFlo, Zoetis). Dogs were positioned in left lateral recumbency and intermittent positive pressure ventilation was maintained throughout using a mechanical ventilator. Image acquisition was performed during transient apnoea, typically induced following a brief period of controlled hyperventilation. Intravenous fluid therapy (at maintenance requirements) was provided with isotonic compound sodium lactate (Aquapharm11, Animalcare) during the procedure. A MRI compatible vectorcardiogram gating device was used to derive electromotive signals in orthogonal spatial planes, to provide R peak triggering for retrospective gating.

### The CMR protocol is summarised in figure 1

#### Standard imaging sequences

Standard imaging sequences were first performed for cine image acquisition using a gradient echo pulse sequence with balanced steady-state free precession (bright-blood imaging). A detailed description is provided in the supplementary information. Images acquired included 2-, 3- and 4-chamber vertical long axis cine images and a series of short axis images of the LV apex of the LV to the dorsal aspect of the LA. Contiguous 5mm slices were acquired, with no interslice gap and 30 frames per cardiac cycle.

**Figure 1:**
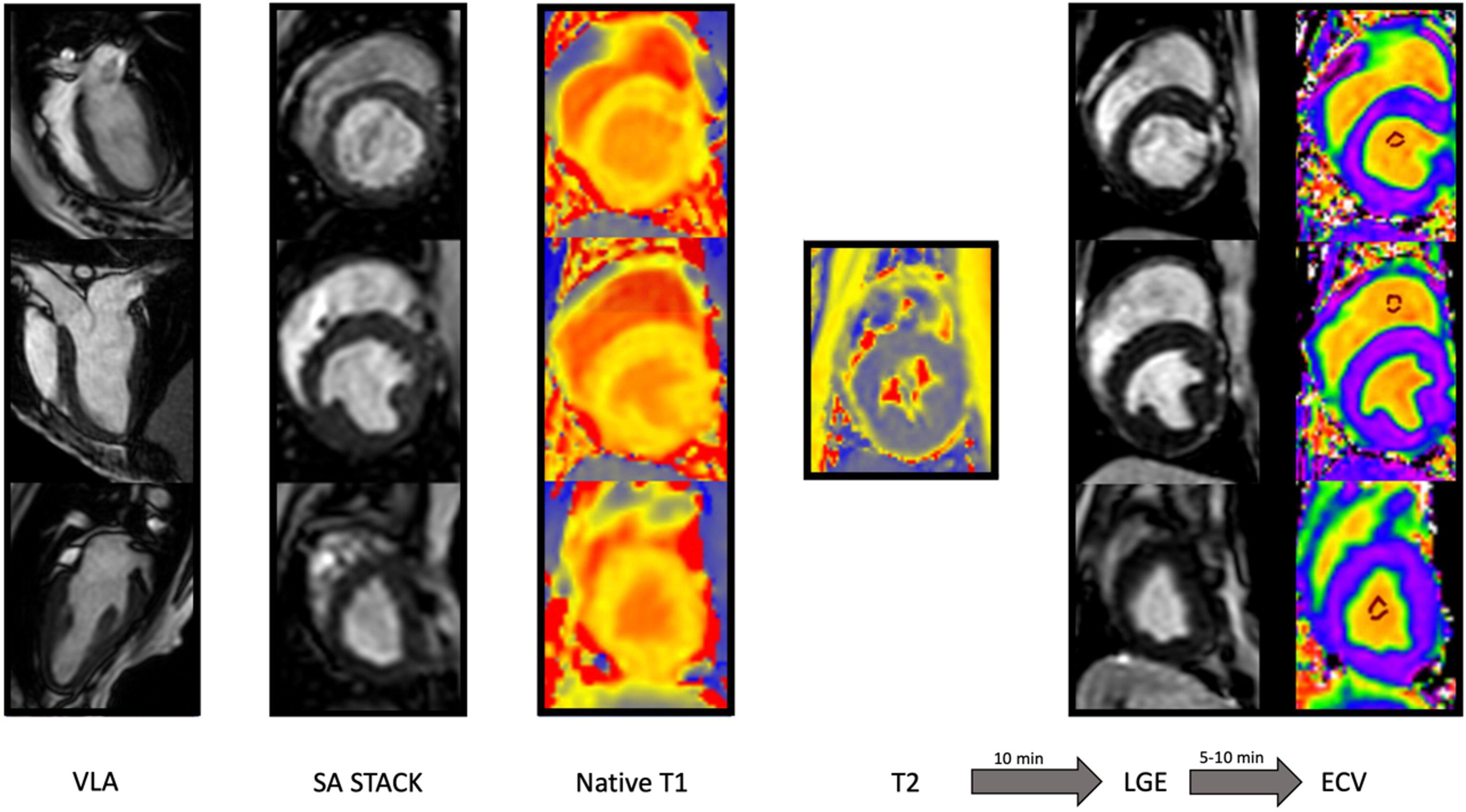
Graphical representation of the cardiac magnetic resonance imaging protocol in dogs aged between 15 and 36 months of age. Vertical long axis images were acquired to prescribe short axis imaging of the left ventricle (LV). Native Tl maps were acquired from the basal, midventricular and apical LV followed by acquisition of midventricular T2 maps. Late gadolinium enhancement imaging was <u>performed</u> 10-15 minutes after injection of 0.2mg/kg gadolinium chelate. Post-contrast Tl maps were collected 15-20 minutes after contrast injection. Post contrast studies were acquired from the same LV levels as for Tl mapping. Extracellular volume was calculated from the combined results of native and post contrast Tl imaging. See text for detailed description. Abbreviations: VLA; vertical long axis. SA; short axis. LGE; late gadolinium enhancement. ECV; extracellular volume.

#### Late gadolinium enhancement imaging

Late gadolinium enhancement imaging was performed between 6 and 12 minutes after intravenous administration of 0.2mg/kg gadolinium (Gd-DTPA (Magnevist)). Images were obtained using inversions recovery prepared turbo fast echo pulse sequences, with optimal inversion time based on visual inspection of the Look-Locker sequence as the time taken effectively to null the normal myocardium. Images were acquired in the 2-, 3- and 4-chamber axes VLA and in short axis at the base, mid and apical myocardial segments. Additional PSIR images were obtained at the same short axis levels to ensure optimal imaging of the ventricular myocardium.

#### Parametric mapping studies

T1 parametric mapping studies were performed using breath-held modified Look-Locker inversion recovery (MOLLI) sequences of the short axis LV myocardium at identical imaging planes to those obtained for LGE imaging. Native studies were acquired immediately prior to contrast administration using ECG-triggered images during diastole. T1 mapping was repeated 15 to 20 minutes after administration of contrast, as detailed above, after LGE imaging. Native T1 maps were acquired using a 5s(3s)3s MOLLI scheme to reduce confounding effects of heart rate on T1 values. Contrast enhanced studies used a 4s(1s)3s(1s)2s scheme, accounting for anticipated shortening of post-contrast T1 times. T2 maps were acquired only from the short axis midventricular LV myocardium using gradient spin echo sequences (GraSE) with breath-holding. Typical imaging acquisition parameters are provided in Table 1. The haematocrit for each dog was obtained from a blood sample drawn immediately prior to induction of general anaesthesia.

**Table 1:** Cardiac magnetic resonance imaging: Typical image acquisition parameters. Philips 1.5T Intera MRI protocols. TE: Echo time. TR: Repetition Time. B-FFE: balanced Fast Field echo (balanced steady state free precession imaging). GRASE: GRadient and Spin Echo sequence for T2 mapping. IR TFE: inversion recovery with Turbo Fast gradient Echo imaging. FA flip angle.

| Sequence | TE (ms) | TR (ms) | Slice thickness (mm) | No. of slices | FA (°) | Field of view (mm<sup>2</sup>) | Matrix | Voxel | Approx. scan time (min) |
| --- | --- | --- | --- | --- | --- | --- | --- | --- | --- |
| **B-FFE** | 2.2 | 3.2 | 5 | 16 | 60 | 137x137x80 | 80x77 | 1.7x1.8x5 | 2.31 |
| **GRASE** | n*11 | 741 | 10 | 1 | 90 | 300x300x10 | 152x140 | 1.9x2.1x10 | 0.09 |
| **MOLLI T1 native** | 2.8 | 1.3 | 10 | 1 | 35 | 300x300x10 | 132x108 | 2.3x2.75x10 | 0.11 |
| **IR-TFE** | 3.5 | 7.1 |  | 12 | 20 | 140x140x60 | 116x102 | 1.2x1.4x5.0 | 2.15 |
| **MOLLI T1 enhanced** | 2.8 | 1.3 | 10 | 1 | 35 | 300x300x10 | 132x108 | 2.3x2.75x10 | 0.12 |

#### CMR post processing

All images were reviewed by the same veterinary cardiologist with 10 years of experience performing canine CMR. Images were first manually reviewed using the Osirix/Horos platform for evidence of bright signal intensity associated with LGE. The presence or absence of LGE accumulation was assessed for each myocardial segment, extrapolating from human nomenclature from the American Heart Association standard 16 segment model.[73] Enhancement had to be present in two imaging planes to be considered genuine, taking care to avoid false inclusion of epicardial fat or intramyocardial vasculature as LGE lesions. Dogs that showed at least one segment with LGE were considered LGE positive.

Parametric mapping data were reviewed using the Qmaps software (Medis Medical Imaging). T1 mapping data were acquired following manual contouring of the LV myocardium from native T1 maps, including the papillary muscles in the adjacent segments. Epicardial and endocardial borders were drawn such that they excluded approximately 10% of the outer- and inner-most myocardium to avoid inadvertent inclusion of the left and right ventricular blood pools and epicardial fat and minimise potential partial volume averaging errors.[40] Regions of LGE were included. Care was taken to avoid basal segments that included the LV outflow tract. Where this inadvertently occurred, those segments were excluded from analysis. Native and post contrast maps were excluded if there were artefacts caused by respiratory motion (sub-optimal breath-holding), evidence of significant banding artefacts, myocardial motion during image acquisition or based on visual assessment of goodness of fit maps during post processing. The LV myocardium of each slice was automatically divided into 6 segments of equal area, following manual selection of the inferior RV septal insertion point. The average T1 value for each ventricular level (base, mid-ventricle, or apex) was calculated as the average value of the 6 segments at that level.[34] The global T1 value for the LV myocardium was calculated as the average T1 values of the basal and midventricular levels. Septal and FW diameter and myocardial area were measured from the same native T1 maps using the Osirix/Horos platform. The heart rate at the time of native and post contrast mapping studies was recorded from the image metadata. Extracellular volume was calculated using the native and post contrast T1 values for each segment and the dog haematocrit using the following equation. [18]

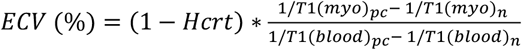

Where Hcrt = haematocrit, T1(myo) = T1 time in the region of myocardium sampled, T1(blood) is T1 time of the sampled blood pool, pc = post contract administration, n = native (prior to contrast administration).

As for native T1, the average ECV value was calculated from the 6 segmental values at each level and the global ECV value calculated as the average ECV value from the base and midventricular levels.

### Postmortem characterisation of myocardial lesions with histopathology

Hearts were excised from all eight DE50-MD dogs immediately after euthanasia, flushed to remove blood clots and fixed whole in 10% neutral buffered formalin. After fixation, the LV from each heart was sectioned in short axis, transverse sections from base to apex. Basal, midventricular and apical sections were selected to approximate the imaging planes obtained during CMR assessments and paraffin-processed to obtain slides of the basal, midventricular and apical LV. Slides were stained routinely with haematoxylin and eosin (H&E), (for 8 hearts) Picrosirius Red (for 7 hearts) and /or Masson’s trichrome (for 8 hearts). All slides were reviewed by a single, American College of Veterinary Pathologists Board-Certified veterinary pathologist, with experience in laboratory animal pathology (including beagles and the *mdx* DMD model). The pathologist was blinded to the results of CMR imaging studies and dog age. Sections were reviewed for recognised histopathological lesions described in the myocardium of DMD patients and canine models of DMD, (juvenile DE50-MD dogs)[63]. Specifically, these were the presence of accumulations of mature adipoctyes, termed “fatty infiltration”, fibrous scarring, acute necrosis, inflammation, oedema, mineralisation, vascular wall abnormalities and Purkinje fibre vacuolation[63]. To avoid overinterpretation of processing artefacts, variability due to plane of section and incidental “background” lesions recognised in healthy canine myocardial specimens, strict criteria for vascular wall abnormalities were used: specifically, there had to be evidence of medial necrosis, disruption of the internal elastic lamina, significant intimal hypertrophy and/or disorganisation of the tunica media with clear expansion of the extracellular matrix.[74, 75] Likewise, oedema was only considered present when increased white space was seen to expand the extracelluar matrix between remaining myocytes in the presence of dilated lymphatics and typically inflammatory infiltrate. Fibrosis was defined by the presence of streams of mature collagen that stained positive on Masson’s trichrome (blue) and picrosirius red (red) with scattered fibroblasts present. Fibrosis and fatty infiltration were awarded a semi-quantitative severity score according to the number of wall segments involved and proportion of the wall layers affected, (i.e., single wall layer versus transmural distribution) (figure 2).

**Figure 2:**
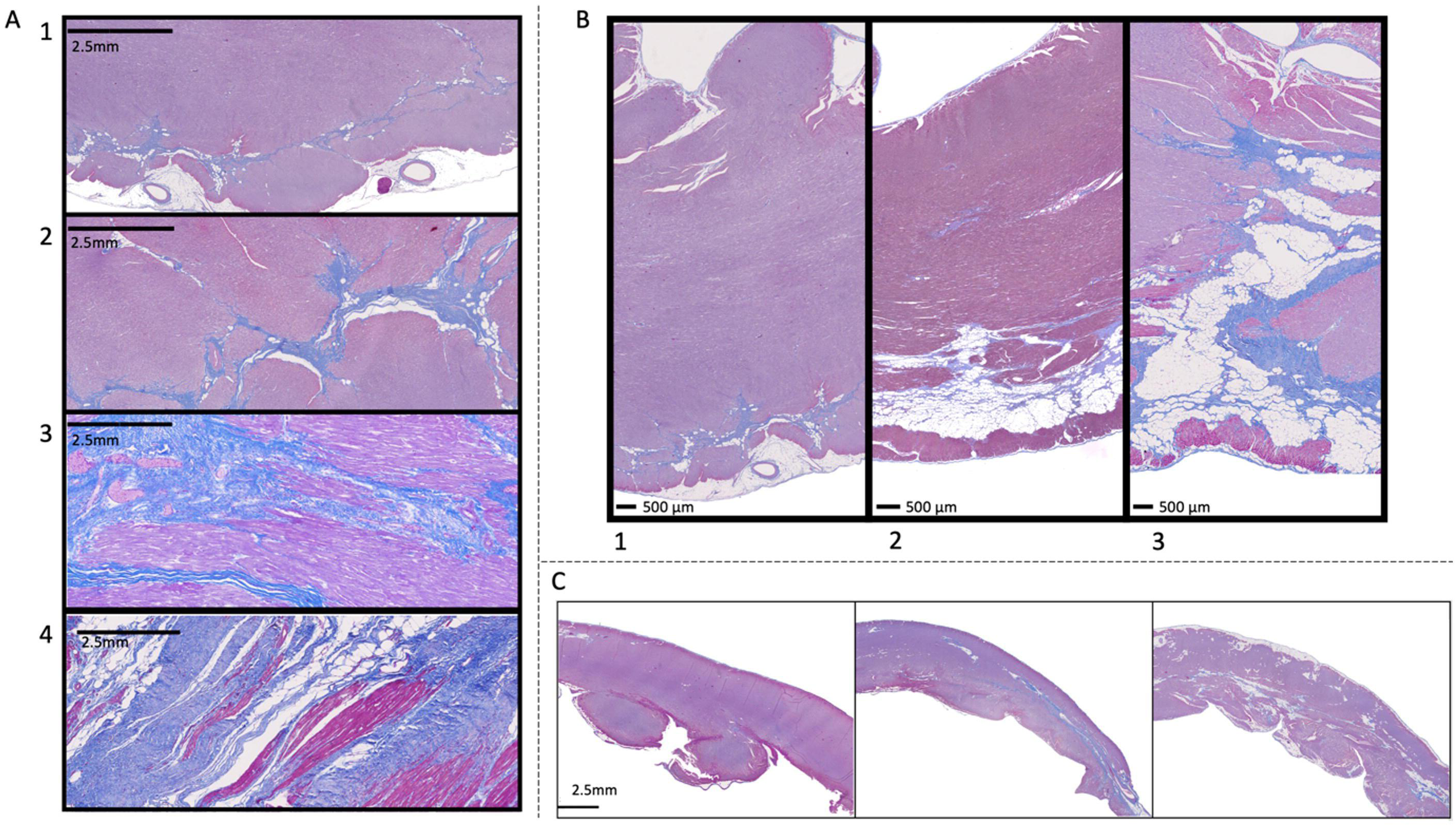
Images depicting regions of the left (A, B) and Right (C) ventricular myocardium stained with Masson’s trichrome. Myofibres stain magenta, whilst collagen fibres stain blue. Figures A and B demonstrate grading system for fibrosis (A) and (B) fatty/adipocyte infiltration (non-staining, vacuolated cells). The score grades were defined as follows: 0: No pathology present. 1: Minimal: Small focal or multifocal lesions. 2: Mild: Single extensive lesion noted but remainder of myocardium unremarkable, or multifocal smaller lesions. 3: Moderate: Multifocally extensive lesions, in single myocardial layer. 4: Marked: As moderate but becoming focally transmural. (5: Severe -not shown: Extensive transmural lesions present - not documented in any DE50-MD dog). Figure C shows sections of right ventricular myocardium from separate DE50-MD dogs demonstrating a spectrum of right ventricular pathological change characterised by mid­myocardial adipocyte and fibrous infiltration of the RV myocardium, increasing in severity from the left to right panel

The semi-quantitative severity scores were defined as follows:

0: No pathology present

1: Minimal: Small focal or multifocal lesions

2: Mild: Single extensive lesion noted but remainder of myocardium unremarkable, or multifocal smaller lesions

3: Moderate: Multifocally extensive lesions, in single myocardial layer

4: Marked: As moderate, but becoming focally transmural

5: Severe: Extensive transmural lesions present

Fibrosis and fatty infiltration scores were combined to generate septal, LV FW, global LV, (combining the septal and FW scores from all sections), and RV pathology scores, (comprising the RV scores from all sections). The presence or absence of fibrofatty replacement for each of the 16 LV myocardial segments (extrapolating from human nomenclature from the American Heart Association standard 16 segment model)[73] was reviewed by the primary investigator and scored as present (segment score 1) or absent (segment score 0).

### Statistical analysis

Statistical analysis was performed using commercially available software (Prism 9.4.1, Graphpad software and SPSS statistics, V28.0/1. IBM). Statistical significance was set at p<0.05. Echocardiographic measures of LV and RV function were compared within genotypes in dogs aged 15 months and those aged 27 to 30 months, (using the oldest time point available), using paired t-tests for parametric data and Wilcoxon matched-pairs signed rank test for nonparametric data. Differences in repeated measures for blood borne biomarkers, (NT-proBNP and cTnI), conventional transthoracic echocardiography and CMR parameters were analysed using a linear mixed model (LMM) with a compound symmetry correlation structure, with age and disease status (genotype), and their interaction entered as fixed effects. The relationship between LGE status and cTnI concentration was evaluated in the same way. Pearson’s correlation coefficient analyses were performed to explore the relationship between predicted physiological/anatomic confounding variables and global native T1 data, both in WT and affected dogs.

The influence of confounding variables on T1 values and their interaction with genotype status were evaluated through inclusion as covariates in LMM analysis with Fisher’s LSD post-hoc comparisons. Genotype, age and their interaction were included as fixed effects and dog identification number as a random effect. Covariates were removed in backwards, step-wise iterations, beginning with the interactions between fixed effects and genotype, but retaining genotype and age and their interaction in the model, to evaluate the effect of repeated measures. Finally, the relationship between global and regional (septal or FW) LV functional parameters, ECG findings, cTnI and results of parametric mapping (average T1, ECV and mid ventricular T2) and LGE analysis from DE50- MD dogs were explored using Pearson’s correlation coefficient. Pearson’s r and p-values are reported.

For all LMM, normality of the computed model residuals of the final model was assessed through visual assessment of histograms and quantile-quantile plots, and finally using the Kolmogorov-Smirnov test for normality. If residuals did not assume a normal distribution, the LMM analysis was repeated following log transformation of the dependent variable. The relationship between the number of LGE segments documented in the final CMR exam and the LV pathology score, and with the number of segments demonstrating fibrous or fibrofatty myocardial replacement was explore using univariable linear regression.

## Results

Results of basic structural CMR and TTE assessments for the study population are summarised in Table 2. Four dogs underwent early euthanasia due to progression of their skeletal muscle phenotype to a humane end point stipulated in the project Home Office licence. One DE50-MD dog was euthanised at 21 months, two dogs at 27 months and one dog at 33 months. In all cases, euthanasia was performed due to dysphagia, which is a common complication in other dystrophin-deficient canine models secondary to reduced jaw opening, macroglossia and pharyngo-oesophageal dysfunction. [76, 77] No other humane endpoint was reached. The dogs euthanised at 21 and 33 months of age respectively underwent additional CMR studies immediately prior to euthanasia to allow comparison between results of histopathology and LGE imaging. A preceding CMR assessment was not possible for either dog that underwent euthanasia at 27 months of age and so there was an interval of 66 and 83 days

**Table 2:**
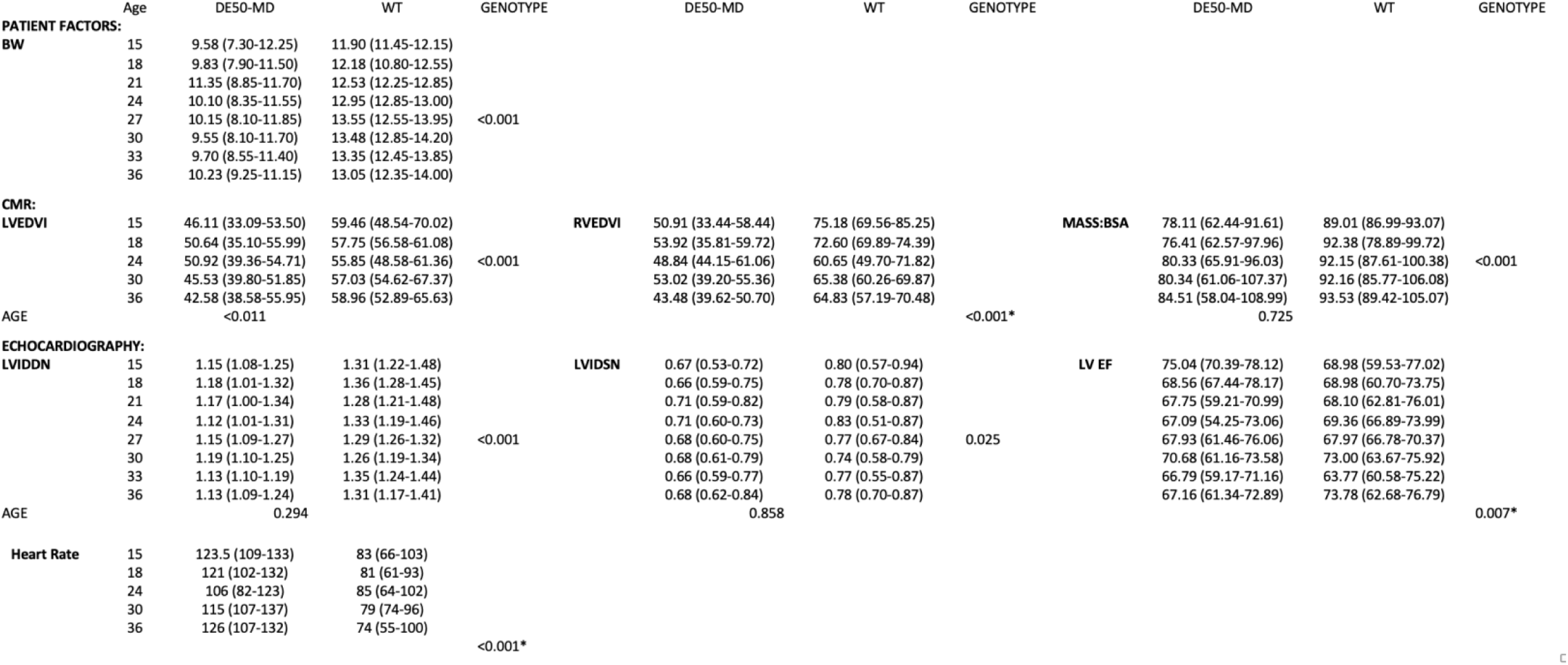
Linear mixed models analysis to explore the effect of genotype and age on structural and functional assessments using cardiac magnetic resonance imaging and conscious transthoracic echocardiography. Abbreviations: WT; Wild type control dogs. BW; Bodyweight. CMR; cardiac magnetic resonance imaging. LVEDVI; Left ventricular end diastolic volume indexed to body surface area. RVEDVI; Right ventricular end diastolic volume indexed to body surface area. LVIDDN; Weight normalised LV diastolic internal diameter. LVIDSN; Weight normalised LV systolic internal diameter. LV EF; Left ventricular ejection fraction. Asterisk signifies significant interaction between age and genotype.

### Cardiac dimensions

DE50-MD dogs had smaller weight-adjusted LV systolic and diastolic diameters, (measured using echocardiography) than WT dogs (p<0.001 and p=0.025 respectively), with no influence of age, (p=0.294 and p=0.858 respectively; Table 2). Left ventricular end diastolic and end systolic volumes (indexed to body surface area) and LVMi derived using CMRI were also smaller in DE50-MD dogs (p<0.001 for all parameters) without a significant influence of age (p=1.00, 0.470 and 0.725 respectively).

### Decline in left and right ventricular systolic function in DE50-MD dogs

Echocardiographic data were available for 5 WT dogs at both 15 and 30 months of age, 7 DE50-MD dogs at 15 months of age, 2 at 27 months of age and 5 at 30 months of age. WT dogs were not evaluated using STE or TDI due to poor tolerance for lateral recumbency. Likewise, left apical views could not be consistently obtained to permit meaningful evaluation of TAPSE in WT dogs.

No dog developed overt systolic dysfunction (EF<55%) during the study period. However, left ventricular EF significantly declined between 15 and 27-30 months in DE50-MD (p=0.003) dogs (figure 3). In contrast, in the WT dogs, EF trended to increase though this did not reach statistical significance (p=0.06). Average peak circumferential systolic strain also declined, (became less negative) in the DE50-MD dogs between 15 and 27-30 months of age, (p=0.006) but SR parameters did not significantly change over the same period, (SSR p=0.460. ESR p=0.827). Septal, (but not anterolateral wall, (p=0.333)) longitudinal velocity also declined in the DE50- MD dogs, (p=0.016, figure 3). Fractional shortening did not change significantly in either group, (DE50-MD dogs, p=0.301. WT dogs p=0.536).

**Figure 3.**
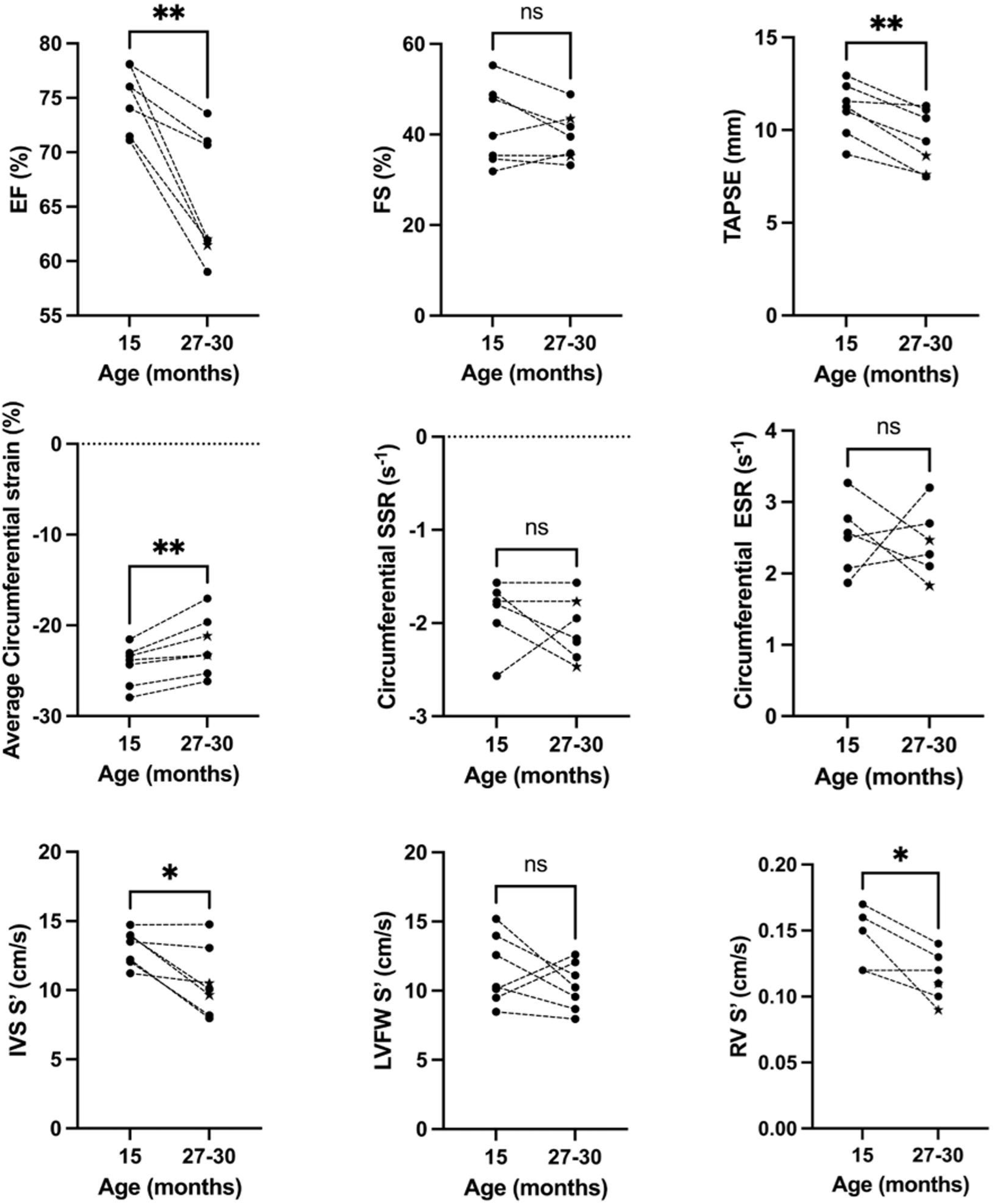
Change in echocardiographic functional parameters between 15 and 27-30 months of age in DESO-MD dogs. Stacked data points indicate results for individual dogs. Individuals represented by a star in the 27-30 group are dogs aged 27 months. Abbreviations: EF; Ejection Fraction. FS; Fractional shortening. SSR; systolic strain rate. ESR; Early diastolic strain rate. IVS S’; Systolic longitudinal velocity of the basal interventricular septum. LVFW S’; Systolic longitudinal velocity of the basal anterolateral (i.e. left ventricular free) wall. RV S’; Systolic longitudinal velocity of the basal Right ventricular wall. ns denotes a non-significance difference between paired values. ** p<0.01 * p<0.05

#### Blood-borne biomarkers

Cardiac troponin I values were within the healthy reference range in all WT dogs at all time points. DE50-MD dogs had higher cTnI than WT dogs at all time points (figure 4). Values exceeding the healthy reference range were obtained in 5/8 DE50-MD dogs during the study period (range 75-25606pg/ml). In three DE50-MD dogs, transient severe elevations in cTnI values (>450pg/ml) were documented at 18 months of age (figure 4) but were not sustained in longitudinal assessment where later timepoints were available (in 2/3 dogs). All other values were below 150pg/ml. Although a significant effect of genotype on cTnI was identified in LMM analysis, a normal distribution of the LMM model residual values could not be obtained, despite attempts to transform the raw data and therefore outlier values were removed. Once removed, cTnI remained significantly higher in DE50-MD dogs (p<0.0001) and this relationship did not change with age (p=0.097). The results of NT-proBNP analysis mirrored that reported in young DE50-MD dogs,[63] such that DE50-MD dogs had significantly lower plasma NT-proBNP than age-matched WT dogs (see supplementary figure 1; genotype p=0.009, age p=0.206). Of the DE50-MD samples taken, 19/23 (83%) were below the lower limit of detection of the assay (250pmol/l), compared to only 5/23 (22%) of WT samples.

**Figure 4.**
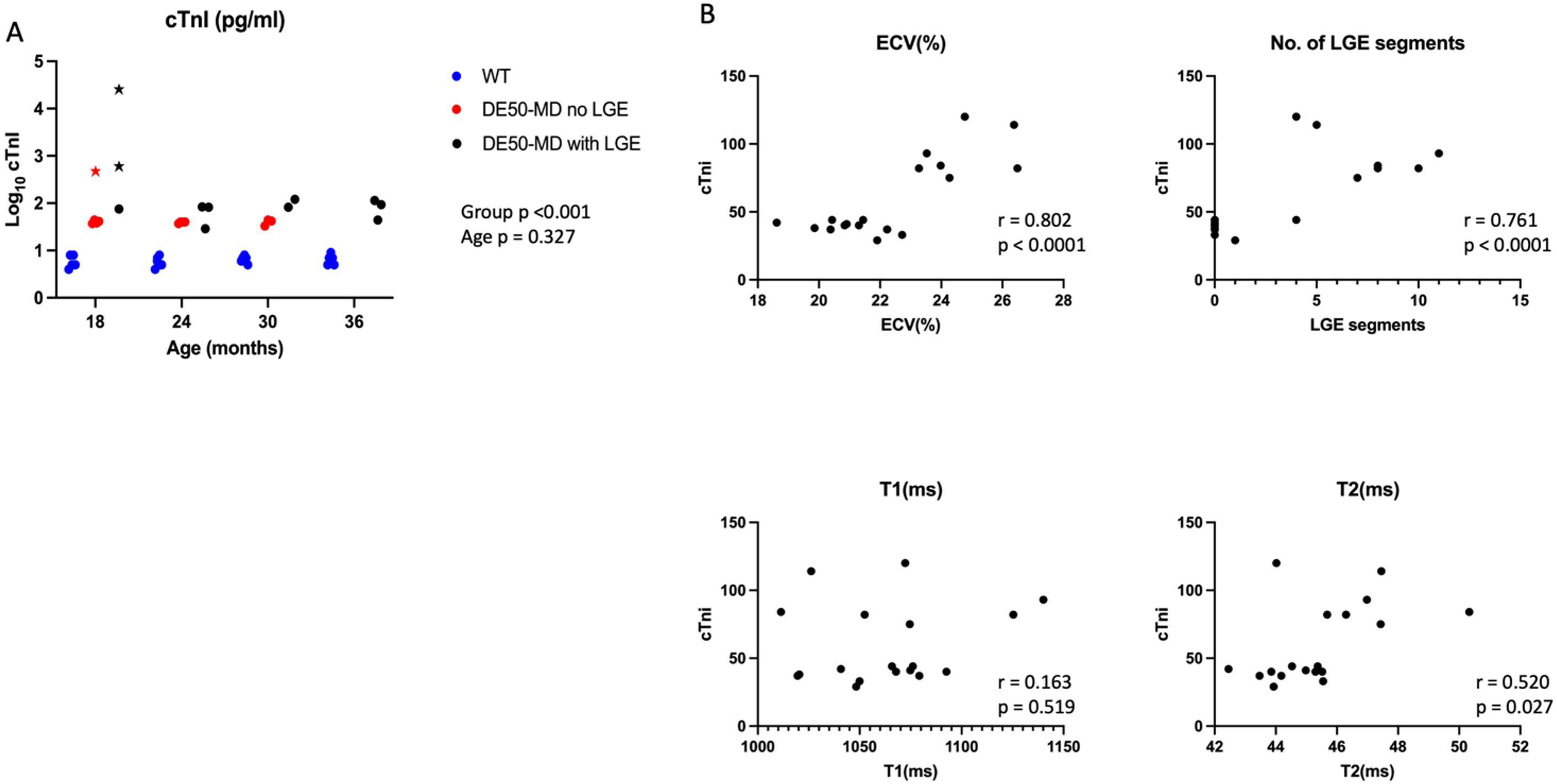
Association between cTnI concentration and age and selected cMRI parameters. (A) cTnl with age in WT dogs (blue, n=6) and DE50-MD dogs, (n=8) with (red) and without (black) LGE aged 18 to 36 months. Each dot represents an individual dog. Those represented by a star had severe transient elevation in serum cTnI at 18 months of age, considered to be outliers in the clinical data. B) Relationship between serum cTnI and parameters derived from cMRI from DE50-MD dogs aged 18 to 36 months. The outlier data points have been removed. Abbreviations: cTnI; cardiac troponin I. LGE; late gadolinium enhancement. ECV; extracellular volume. T2; Average midventricular T2 time. Tl; Average Tl time. A moderate linear association was demonstrated between average T2 and cTnI <u>(r=0.520</u> p=0.0027) and strong linear association was identified for ECV and number of LGE segments with cTnl <u>(r=0.802</u> and <u>r=761</u> both respectively, p<0.0001) There was no association identified between Tl and cTnI.

#### Late Gadolinium Enhancement

LGE was identified in 75%, (6/8) DE50-MD dogs between 15 and 36 months of age, its prevalence increasing with age, (see table 3). In LGE positive dogs, the number of segments with LGE increased with subsequent visits. Enhancement was identified in the outer third of the myocardium (subepicardially) in basal and midventricular LV FW segments, primarily in the antero- and inferolateral wall segments, extending to the inferior or anterior wall segments, (figure 5). LGE was identified in septal segments in 50% (3/6) DE50-MD dogs with LGE. Two of the three dogs had shown LGE accumulation exclusively in FW segments in the CMR assessment 3-6 months prior and septal LGE accompanied FW LGE in all cases. Septal LGE tended to be mid myocardial or located towards the RV endocardial surface. LGE was not identified in any WT dog at any age. cTnI concentrations were significantly higher in DE50-MD dogs with LGE compared to those without, (p<0.001) but results in DE50-MD dogs still significantly exceeded those of WT dogs, even when LGE was not present, (p<0.001) and this relationship did not change with age, (p=0.327, figure 4).

**Figure 5.**
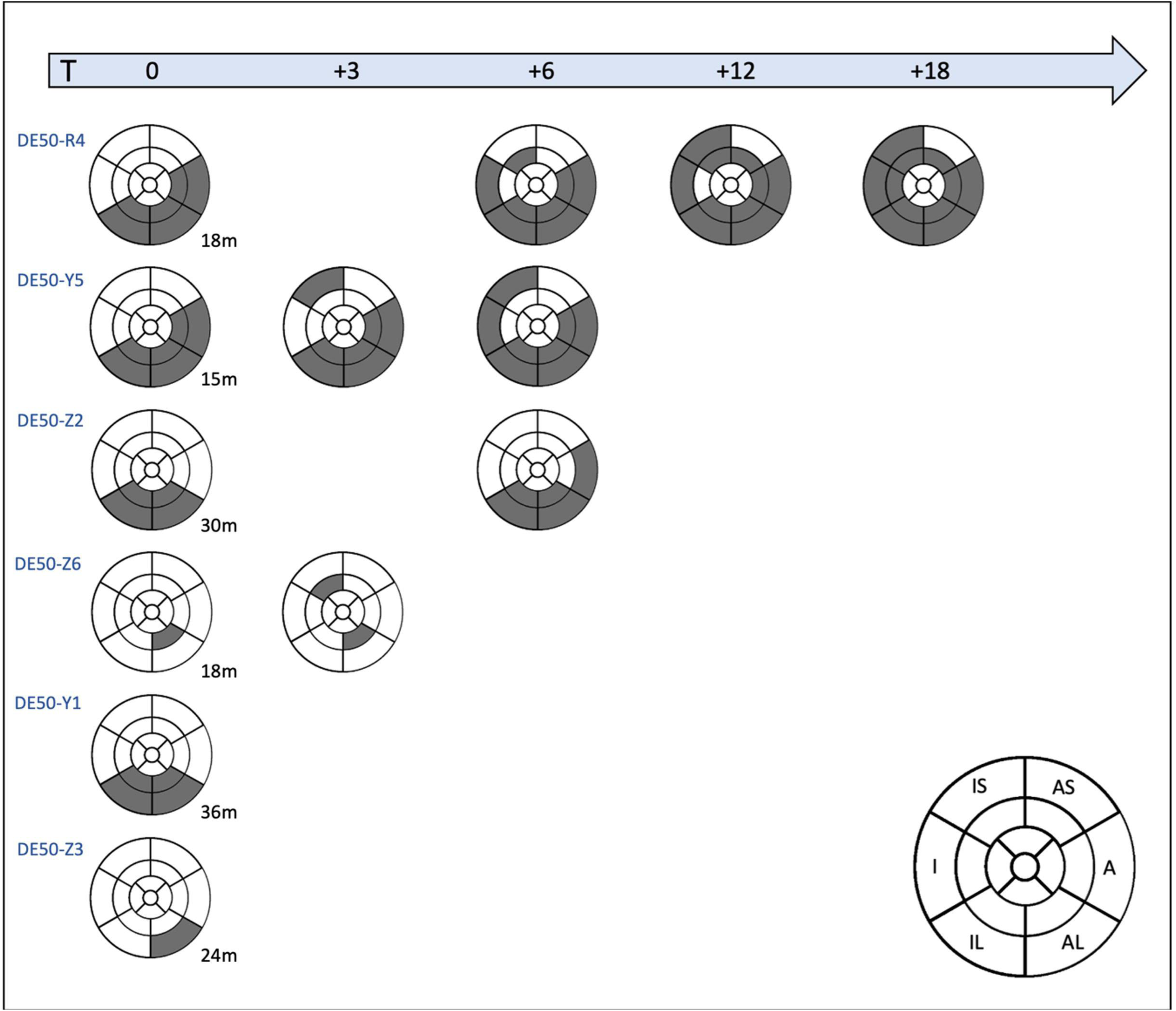
Schematic to demonstrate serial late gadolinium enhancement (LGE) studies in LGE positive DESO-MD dogs, showing the distribution and progression of LGE over time. Each row represents an individual dog, (labelled with dog ID number and the age at which LGE was first identified) while each column represents the time (in months) after first evidence of LGE (Time (T) = 0). Anterolateral and/or inferolateral midventricular wall segment involvement was seen in all LGE positive dogs. Septal involvement was only present when free wall segment involvement was already present.

**Table 3:** Late gadolinium enhancement (LGE) imaging Showing the frequency of the appearance of LGE positive segments in DE50-MD dogs aged between 15 and 36 months. The number of dogs exhibiting LGE increased over time and, for each dog demonstrating LGE, the number of segments that were LGE positive increased over time.

| Age (months): | 15 | 18 | 24 | 30 | 36 |
| --- | --- | --- | --- | --- | --- |
| Dogs with LGE (n) | 1 | 3 | 3 | 2 | 3 |
| Dogs imaged (n) | 8 | 8 | 7 | 5 | 4 |
| Dogs with LGE (%) | 12.50 | 37.50 | 42.86 | 40.00 | 75.00 |
| LGE +ve segments (n) | 6 | 14 | 17 | 14 | 20 |
| Segments imaged (n) | 128 | 128 | 112 | 80 | 64 |
| Segments with LGE (%) | 4.69 | 10.94 | 15.18 | 17.50 | 31.25 |

#### Parametric mapping

Regional T1, T2 and ECV data are summarised in Supplementary table 1.

### Identification of confounding variables: T1 mapping

The mean T1 values of the basal and midventricular segments were similar in within-group analyses (DE50-MD dogs p= 0.721, WT dogs p=0.169). Analysis of apical segments revealed that the average T1 values for the apical segments was significantly higher than that of the midventricular and basal LV segments (p<0.001 for each, for both genotypes). Apical segments were excluded in subsequent analyses.

Pearson’s correlation coefficient analysis was performed to investigate the association between estimated T1 values and potential confounding physiological/patient factors (supplementary figure 2) in dogs aged 15-36 months of age. Results were available for 43, (WT=21, DE50-MD = 23) basal and 60, (WT=30, DE50-MD=30) midventricular slices. A highly significant effect of heart rate was demonstrated on recorded T1 values (DE50- MD 0.501, p=0.001, WT r=0.434, p<0.001). In WT dogs, the T1 value was negatively correlated with myocardial area (r=-0.304 p=0.03) and myocardial thickness (r=-0.370, p=0.008). These associations were not demonstrated in the DE50-MD dogs (r=0.179, p=0.203 and r=0.110, p=0.436 respectively). On evaluation of both the basal and midventricular slices, myocardial area was reduced (p=0.009 and p=0.022 respectively) and the recorded heart rate was increased (p<0.001 for both) in DE50-MD dogs compared WT dogs. Myocardial thickness was similar between genotypes for both ventricular levels (p=0.923 and p=0.475 respectively) (see supplementary figure 3). Given that there was a moderate, significant relationship between heart rate and myocardial area, (r=0.541, p<0.001) but not between heart rate and myocardial thickness (r=0.171, p=0.08), and that myocardial thickness had a greater strength of association than myocardial area with T1 values in WT dogs, myocardial thickness was selected over area as a covariate in LMM.

### Identification of confounding variables: ECV mapping

After exclusion of apical slices, the estimated ECV values of the basal and midventricular LV were explored in LMM, entering myocardial thickness and heart rate as potential covariates, alongside level of sampling, age and genotype. The ECV values required natural logarithmic transformation (Ln) to achieve normality of LMM residuals. Results of the LMM revealed that (Ln)ECV was increased at all ages (age*genotype interaction p=0.007), but that (Ln)ECV was significantly higher (without any interaction with genotype) in the basal slice than the mid ventricular slice (p=0.006). The ECV values from the midventricular and basal slices were therefore averaged to calculate the global ECV for each dog for subsequent analyses.

### Comparison of results of parametric mapping studies between DE50-MD and WT dogs

#### Study Population

Initial studies from WT dogs were performed early in the development of protocols for parametric mapping when midventricular and native T1 acquisitions were prioritised. Due to the lack of native and post contrast T1 imaging from basal segments recorded in WT dogs aged 15 months, estimated values for myocardial T1, T2 and ECV were only compared between genotypes in dogs aged 18-36 months.

Average T1 values were compared between DE50-MD and WT dogs aged 18-36 months in LMMs. To account for the fact that native T1 values from both the midventricular and basal levels from the same dogs were entered separately into LMM, the influence of the ventricular level of sampling was also explored in the LMM, alongside genotype and age, with no significant effect identified. Despite adjustment for myocardial thickness and heart rate, (which remained significant covariates in LMM), native T1 values were significantly higher in DE50-MD dogs from 30 months of age when compared with WT dogs (figure 6).

**Figure 6.**
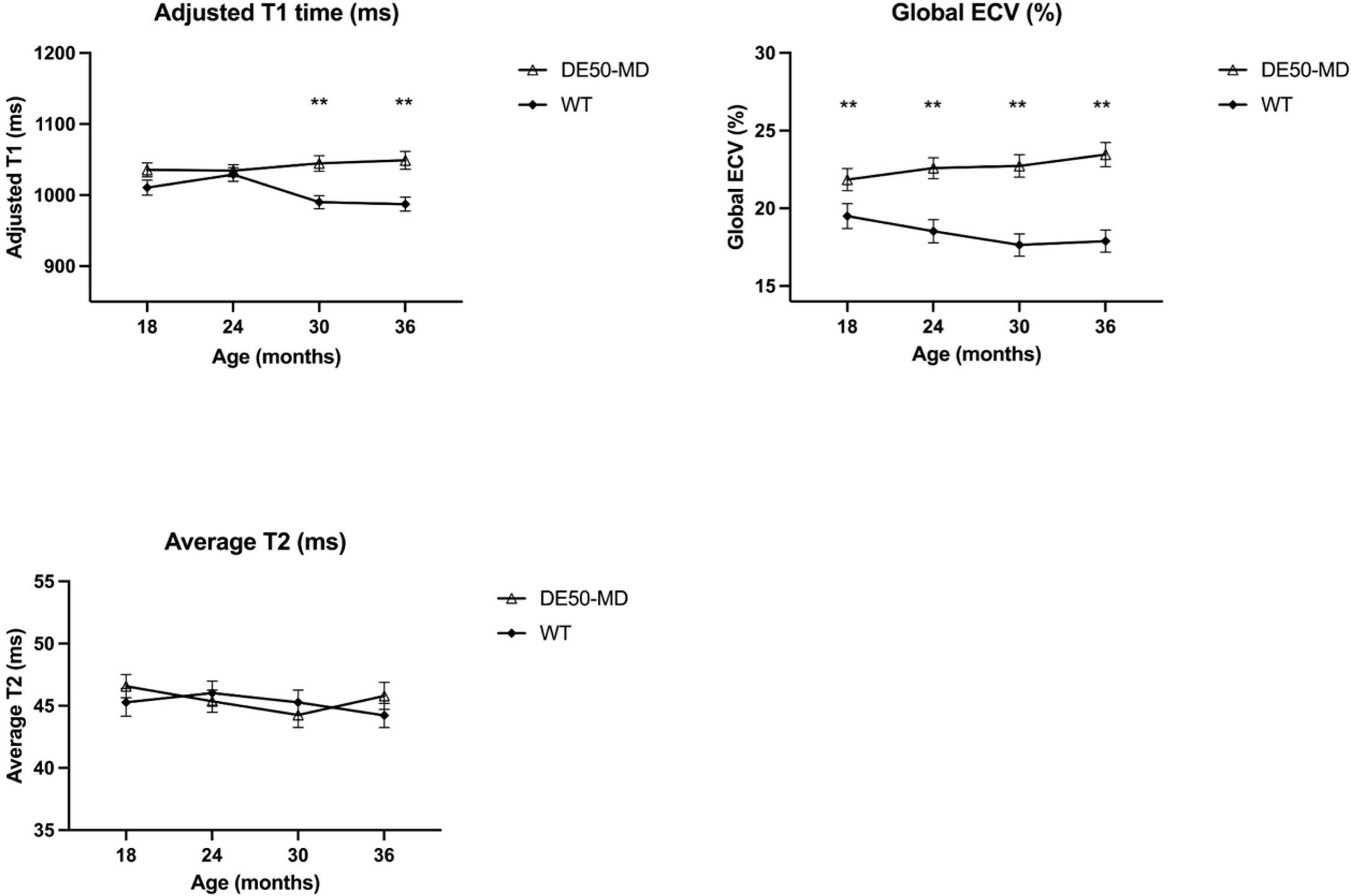
Results of parametric mapping studies in DESO-MD and wild type control (WT) dogs. Complete parametric mapping studies were <u>performed</u> every 6 months between 18 and 36 months of age. Symbols indicate the mean value for each genotype, while error bars represent the standard error of the mean for each genotype at each time point. i) Average Tl times from the base and midventricular myocardium were adjusted according to significant covariates, heart rate and myocardial thickness. A significant difference in adjusted Tl times between genotypes was identified in dogs from 30 months of age (DE50-MD n=4-7. WT n=6)ii) Global extracellular volume (ECV) was significantly higher in DE50-MD dogs compared to WT dogs at all ages studied, with the difference between groups increasing over time (DE50-MD n=4-7. WT n=4-6). iii) No significance was identified in average midventricular T2 times between genotypes (DE50-MD n=4-7. WT n=4-6). ** p<0.001

Global ECV was compared between DE50-MD and WT dogs aged 18 to 36 months of age in LMM analysis. None of the previously identified covariates for native T1 values (myocardial thickness, mid ventricular T2 value or heart rate) remained significant when entered in the LMM. The interaction between genotype and age remained significant (p<0.001). DE50-MD dogs had significantly increased ECV at all ages (p<0.001), the mean difference increasing with age, (mean difference in ECV was 2.3% at 18 months, increasing to 5.6% at 36 months, figure 6). No significant influence of genotype was demonstrated on average midventricular T2 values for DE50-MD dogs and WT dogs (p=0.863) and results did not change with age (p=0.384, figure 6).

#### Relationship between results of CMR tissue characterisation and results of cTnI, electrocardiographic and echocardiographic studies in DE50-MD dogs

After removal of outlier values, cTnI measurements showed a moderate association with T2 values, (r = 0.520, p=0.027) and a strong association with ECV, (0.802, p<0.0001) and number of LGE segments, (r = 0.761, p<0.001) but not native T1 values (r = 0.163, p= 0.519). The relationships between results of parametric mapping and LGE imaging and conscious echocardiographic functional parameters are summarised in table 4.

**Table 4:**
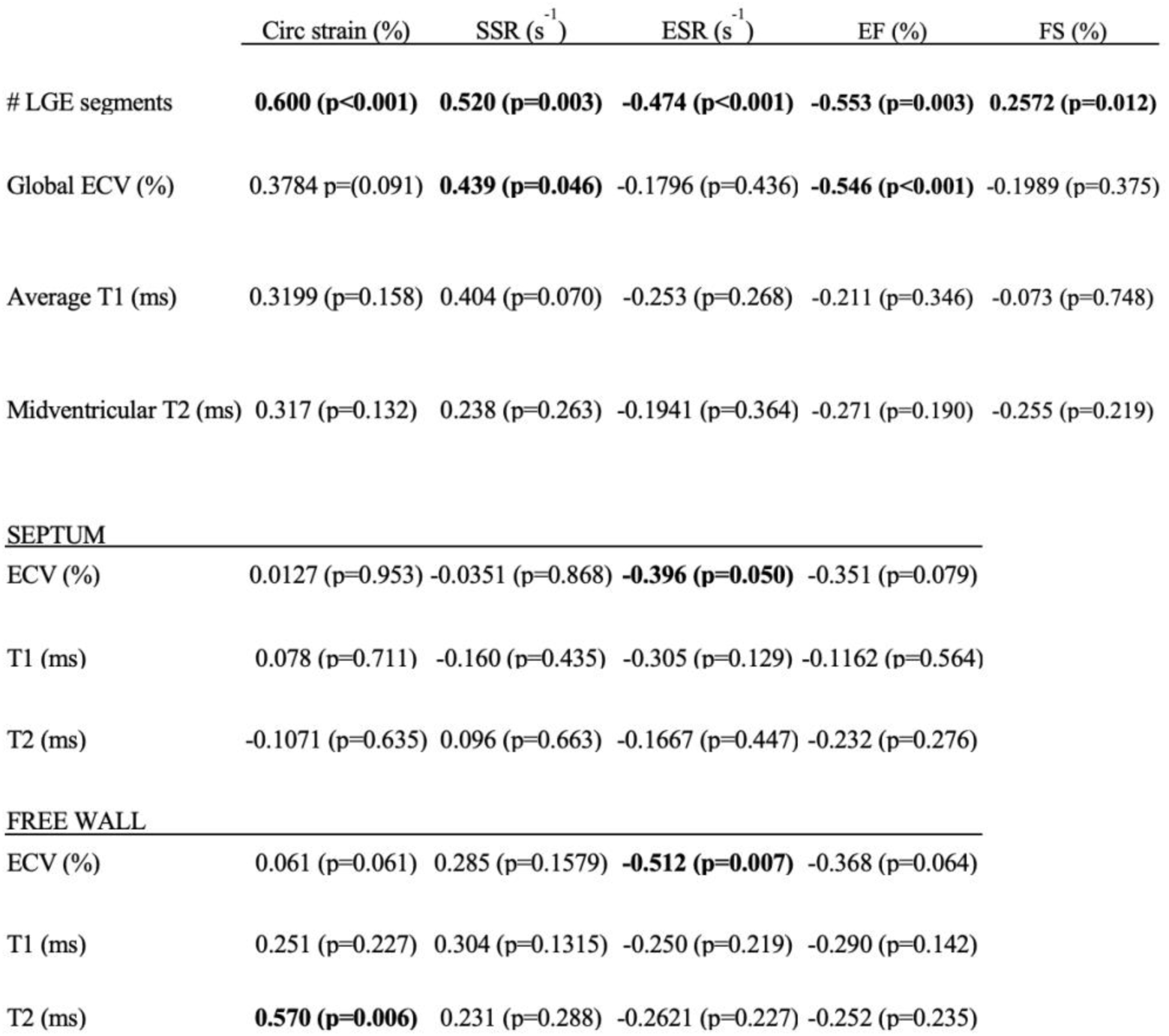
Pearson’s correlation coefficient analyses to demonstrate the relationship between late gadolinium enhancement imaging, parametric mapping and conscious echocardiographic functional assessments. Data is presented as (Pearson’s r value, P value). Abbreviations: Circ strain; peak circumferential strain. SSR; circumferential systolic strain rate. ESR; circumferential early diastolic strain rate. EF; ejection fraction. FS; fractional shortening. # LGE segments; number of myocardial segments with LGE, ECV; extracellular volume. T1; T1 relaxation time. T2; T2 relaxation time

| | Circ strain (%) | SSR ($s^{-1}$) | ESR ($s^{-1}$) | EF (%) | FS (%) |
| --- | --- | --- | --- | --- | --- |
| # LGE segments | **0.600 (p<0.001)** | **0.520 (p=0.003)** | **-0.474 (p<0.001)** | **-0.553 (p=0.003)** | **0.2572 (p=0.012)** |
| Global ECV (%) | 0.3784 p=(0.091) | **0.439 (p=0.046)** | -0.1796 (p=0.436) | **-0.546 (p<0.001)** | -0.1989 (p=0.375) |
| Average T1 (ms) | 0.3199 (p=0.158) | 0.404 (p=0.070) | -0.253 (p=0.268) | -0.211 (p=0.346) | -0.073 (p=0.748) |
| Midventricular T2 (ms) | 0.317 (p=0.132) | 0.238 (p=0.263) | -0.1941 (p=0.364) | -0.271 (p=0.190) | -0.255 (p=0.219) |
| **SEPTUM** |  |  |  |  |  |
| ECV (%) | 0.0127 (p=0.953) | -0.0351 (p=0.868) | **-0.396 (p=0.050)** | -0.351 (p=0.079) |  |
| T1 (ms) | 0.078 (p=0.711) | -0.160 (p=0.435) | -0.305 (p=0.129) | -0.1162 (p=0.564) |  |
| T2 (ms) | -0.1071 (p=0.635) | 0.096 (p=0.663) | -0.1667 (p=0.447) | -0.232 (p=0.276) |  |
| **FREE WALL** |  |  |  |  |  |
| ECV (%) | 0.061 (p=0.061) | 0.285 (p=0.1579) | **-0.512 (p=0.007)** | -0.368 (p=0.064) |  |
| T1 (ms) | 0.251 (p=0.227) | 0.304 (p=0.1315) | -0.250 (p=0.219) | -0.290 (p=0.142) |  |
| T2 (ms) | **0.570 (p=0.006)** | 0.231 (p=0.288) | -0.2621 (p=0.227) | -0.252 (p=0.235) |  |

The number of LGE positive wall segments and global ECV values showed the closest association with LV function. The number of LGE positive segments had a moderate negative correlation with EF, (r=-0.533, p<0.001) and circumferential ESR, (r=-0.474, p<0.001) and a moderate positive correlation with peak circumferential strain, (r=0.600, p<0.001) and SSR (r=0.520, p=0.003) i.e. values became less negative with increasing numbers of LGE positive segments. Global ECV also showed a mild to moderate negative correlation with EF, (r=-0.456, p<0.001) and weak positive association with circumferential SSR (r=0.439, p=0.046). Global T1 and midventricular T2 values did not show any significant correlation with EF or circumferential strain deformation parameters (table 4).

When considering regional parametric mapping parameters, ECV of the LV FW showed a moderate negative correlation with regional circumferential ESR, (r=-0.512, p=0.007). Septal ECV also showed a weak correlation with regional circumferential ESR, but this did not quite reach statistical significance (r=-0.396, p=0.050). Left ventricular FW (FW) T2 values showed a moderate positive correlation with the FW peak circumferential strain (r=0.570, p=0.006) but no other relationships between regional T2 or T1 were identified.

Fragmented ECGs were identified in 7/8 DE50-MD dogs over the study period, the number of leads demonstrating fQRS complexes tending to increase with age in each dog. There was a strong positive correlation between number of leads with fQRS and both the number of LGE segments (r=0.844, p<0.0001), global ECV (r=0.643, p<0.001) and a weaker association with global T1 (r= 0.476, p=0.025) and midventricular T2 times, (r=0.563, p=0.003). The number of leads with fQRS was also moderately negatively correlated with EF, (r=-0.442, p=0.002) and early diastolic circumferential strain rate, (r=-0.350, p=0.020) and strongly correlated with average peak circumferential strain, (r=0.643, p<0.0001) and systolic strain rate (r=0.555, p<0.0001. figure 7).

**Figure 7.**
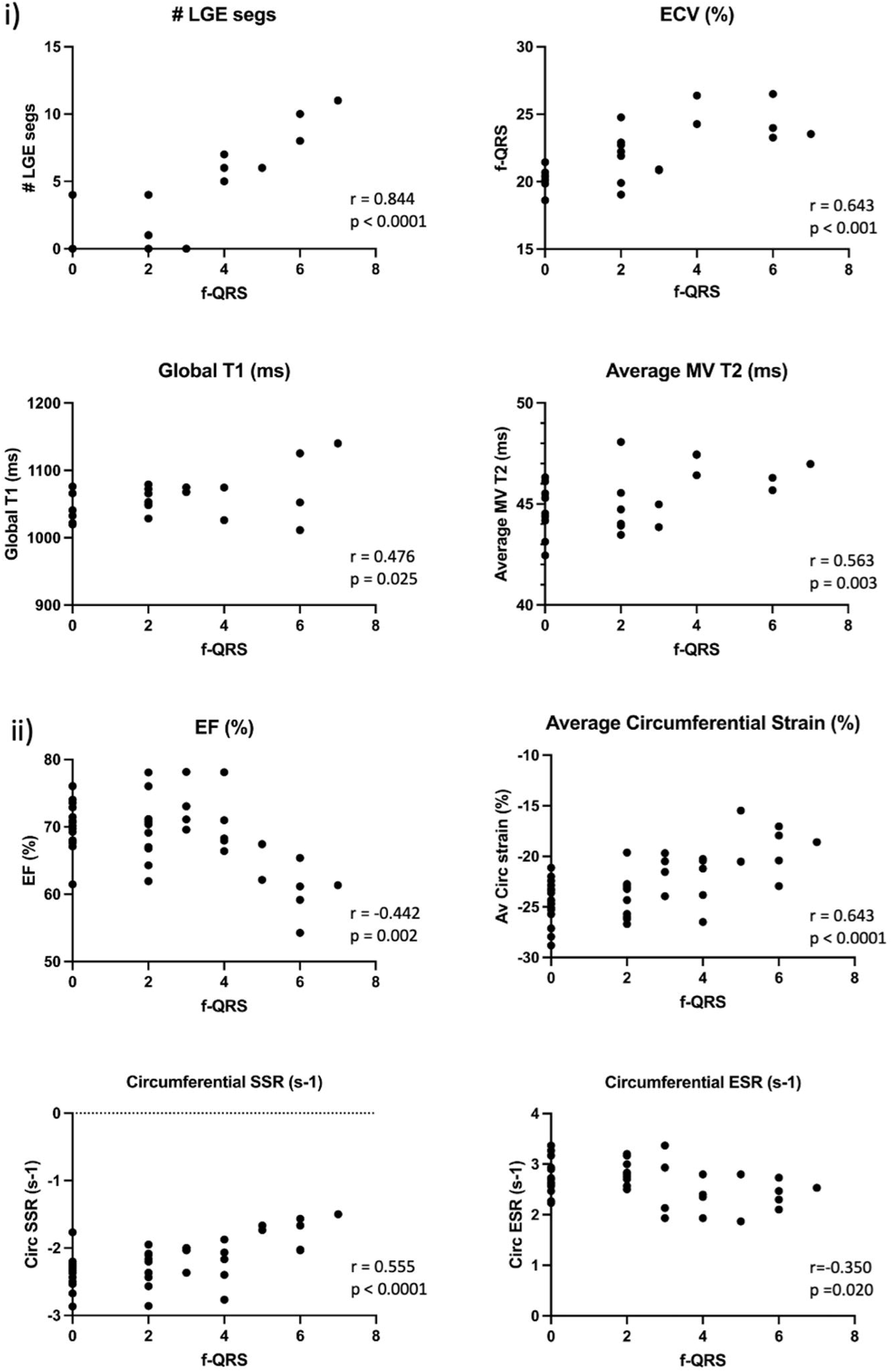
Relationship between the number of leads demonstrating fragmented QRS complexes (f-QRS) and parameters derived from i) cardiac magnetic resonance imaging and ii) echocardiography from DESO-MD dogs aged 15 to 36 months. Abbreviations: # LGE segs; Number of segments demonstrating late gadolinium enhancement. ECV; extracellular volume. MV T2; Average midventricular T2. EF; ejection fraction measured with echocardiography. Circ; circumferential. SSR; systolic strain rate. ESR; early diastolic strain rate

#### Comparison between results of late gadolinium enhancement imaging and histopathology

In agreement with results of antemortem CMR imaging findings, DE50-MD dogs had a spectrum of cardiac histopathological changes. In general, lesions were more prominent in older dogs, however 2/5 dogs aged between 33-36 had minimal pathological change, as demonstrated by low overall LV and RV pathology scores (see table 5).

**Table 5:**
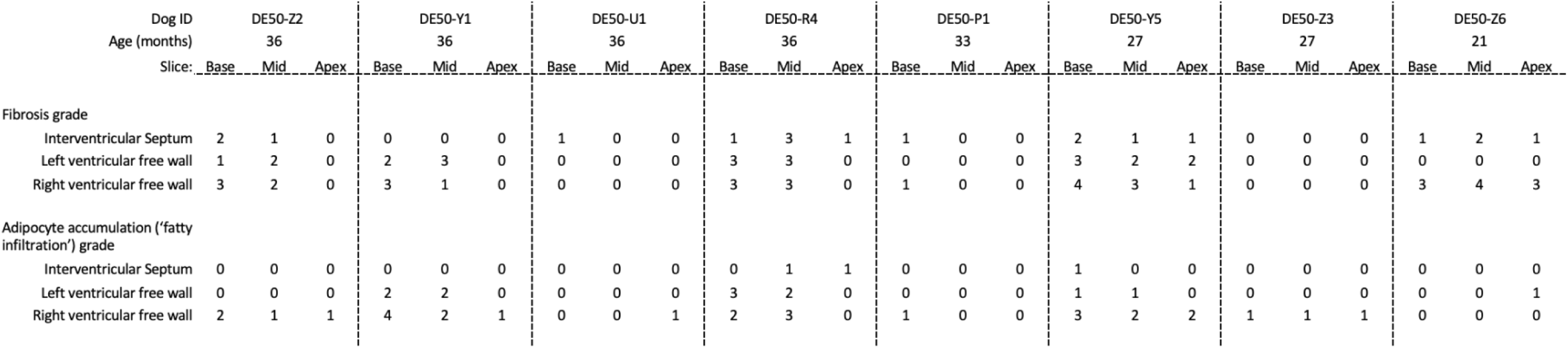
Fibrosis and fatty/adipocyte infiltration scores from the left and right ventricular free walls and septum of individual dogs in basal, mid ventricular and apical sections. The score grades were defined as follows: 0: No pathology present. 1: Minimal: Small focal or multifocal lesions. 2: Mild: Single extensive lesion noted but remainder of myocardium unremarkable, or multifocal smaller lesions. 3: Moderate: Multifocally extensive lesions, in single myocardial layer. 4: Marked: As moderate, but becoming focally transmural. 5: Severe: Extensive transmural lesions present (not documented in any DE50-MD dog).

The principal findings were myocardial fibrosis, with or without accumulation of adipocytes or “fatty infiltration” of the myocardium. Although multiple segments were involved, no individual dog scored higher than grade 3 for fibrous or fatty infiltration lesions in the LV. Overall these pathologies predominately affected basal and midventricular sections, while apical segments tended to be spared. Lesions typically extended in bands between intramural coronary vasculature, in a characteristic circumferential seam within the subepicardium of the left ventricular FW (figure 8). In one dog (aged 36 months), fatty infiltration lesions focally approached a near transmural distribution in the basal FW (figure 2B). Lesions were primarily fibrotic in dogs with lower LV pathology scores, with a greater tendency for fatty infiltration to predominate in dogs with overall higher LV pathology scores. As for results of LGE imaging, septal lesions were typically located within the midmyocardium, or towards the RV aspect. Lesions in the septum tended to be fibrous, while fatty infiltration was rare, such that only minimal, (grade I) fatty infiltration was detected in the basal septum of two dogs. These dogs also had the highest overall fibrosis and fatty replacement pathology scores. Right ventricular involvement was prominent in DE50-MD dogs, (table 5). In the RV, pathology was distributed predominately in the mid-myocardium, appearing as concentric, anastomosing bands of myocardial fibrofatty replacement (figure 2C). The pathology scores of the RV and LV had a strong association in univariable linear regression analysis (R^2^ 0.703, p<0.001, figure 9).

**Figure 8.**
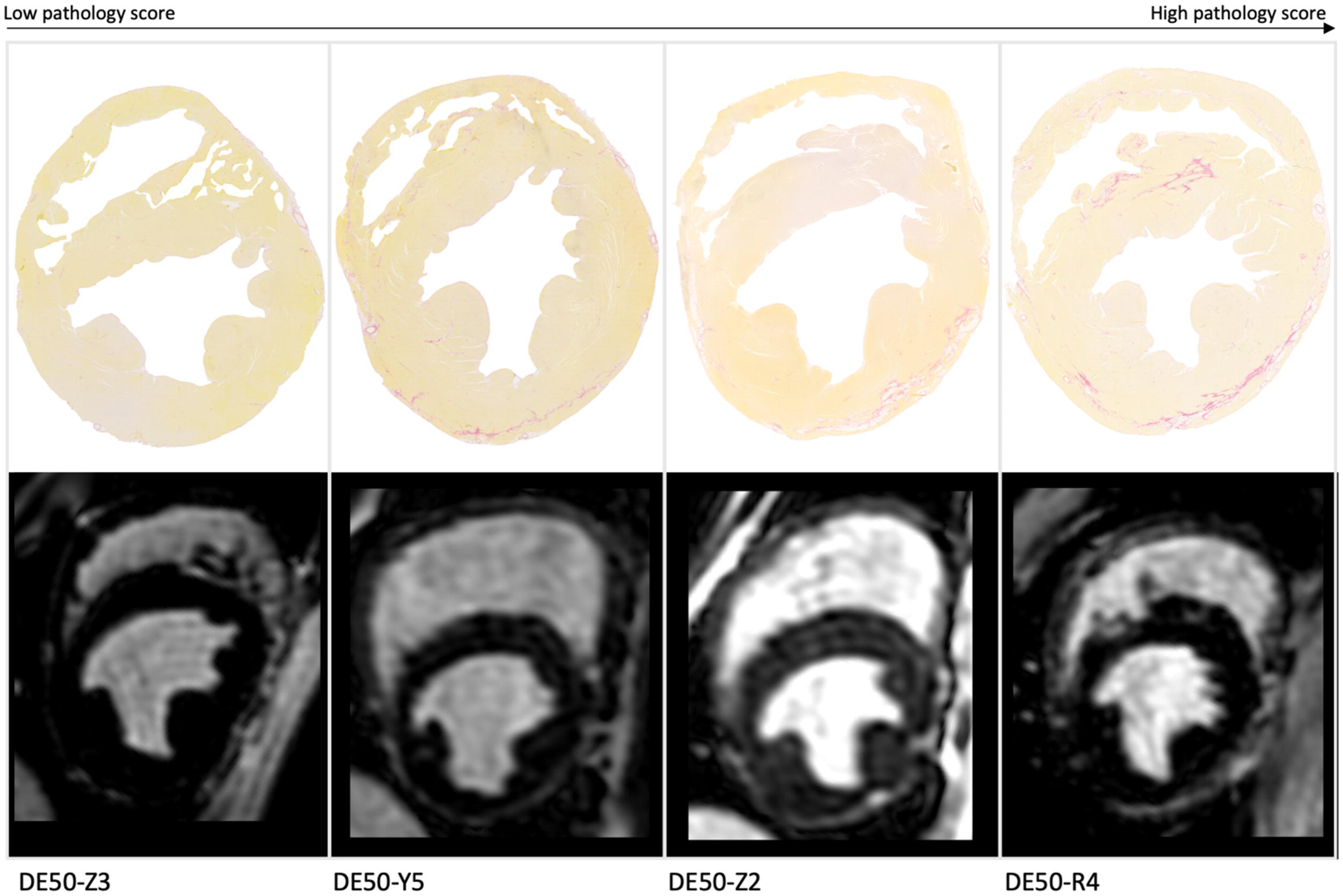
Examples of cross-sectional imaging studies. The upper panel shows histological sections from individual dogs at the midventricular level, stained with picrosirius red. Myocardial cells are stained yellow and collagen is stained red. The lower panel shows the corresponding late gadolinium enhancement (LGE) image from the fmal studies of each dog. Regions of LGE appear as white hyperintense signal next to the nulled (black myocardium) and correspond to regions of myocardial fibrofatty replacement.

**Figure 9:**
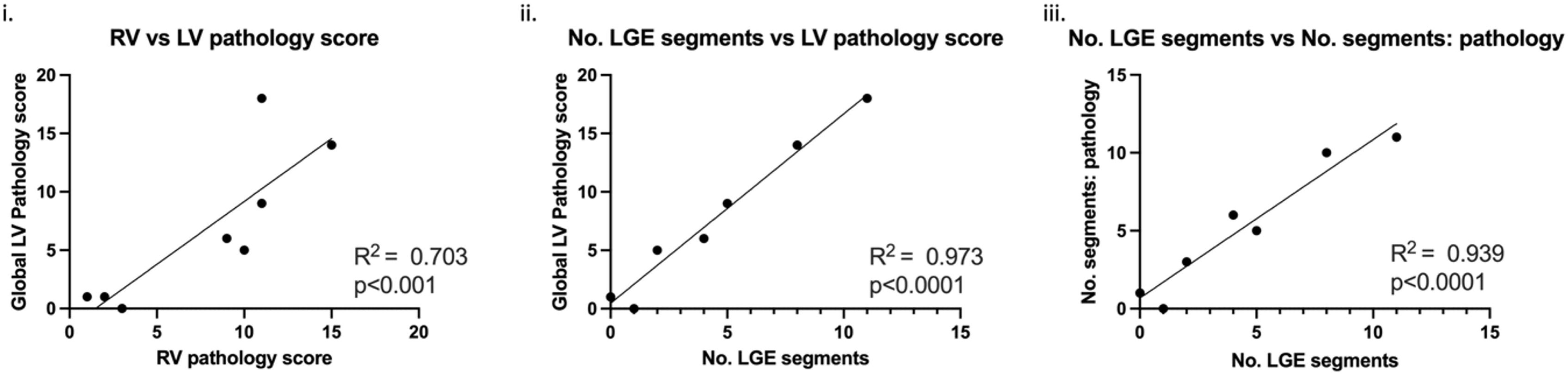
Results of univariable logistic regression analysis to explore the relationship between i. left ventricular (LV) and right ventricular (RV) pathology scores and ii./iii. the relationship between results of histopathological analysis and no. (number) of late gadolinium enhancement (LGE) positive segments detected during cardiac magnetic resonance imaging.

Foci of myocyte necrosis were only identified in one dog - the youngest dog studied. In this dog, the regions of necrosis were accompanied by lymphohistiocytic inflammation without fibrosis (confirmed by absence of collagen straining with Masson’s Trichrome) and were present within the mid and basal septal and RV wall, and focally within the apical anterolateral papillary muscle (figure 10A). The same dog had accompanying myocardial oedema. Overall changes were considered consistent with an acute/recent myocardial insult, likely representing an early lesion prior to onset of fibrotic scarring. These lesions could be identified by LGE accumulation on cross sectional imaging. Lymphohistiocytic inflammatory infiltrates were noted in two additional dogs, which were the dogs with the two highest LV pathology scores (see table 5). Mineralisation was not a feature of the cardiomyopathy in DE50-MD dogs.

Rare, subtle vascular wall changes were noted in the intramural coronary vasculature in 2 dogs (Figure 10B). Changes were characterised by expansion of the tunica media by disorganised extracellular matrix and disruption of the elastic lamina, with possible evidence of oedema. Elsewhere, vessels demonstrated variable tunica media thickness, but resembled those reported in healthy beagles, (and WT dogs in GRMD studies[78]) consistent with the normal variation in the appearance of mural arteries in the canine heart.

**Figure 10:**
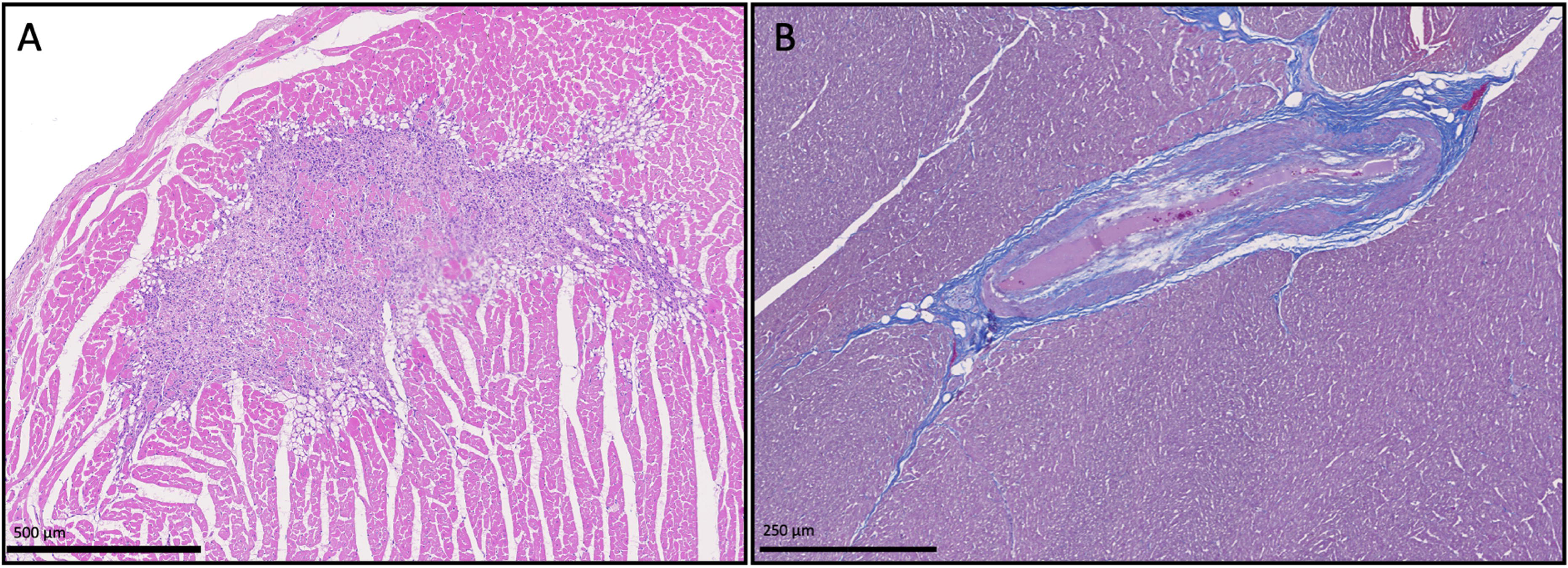
Region of myocardial necrosis (A) and intramural coronary vascular wall changes (B) found in DESO-MD dogs. A) Section of the anterio-lateral papillary muscle of a 20-month-old DE50-MD dog. Haematoxylin and eosin x 2.5 magnification. Image demonstrates a focal region of myocyte necrosis, accompanied by surrounding lymphohistiocytic <u>inflammatory</u> infiltration. B) An intramural arteriole of a 36-month-old DE50-MD dog demonstrating disruption and fibrosis of the tunic media, absence of the elastic lamina and evidence of oedema in Masson’s trichrome.

Regions of LGE mostly corresponded to regions of fibrofatty scarring within the myocardium of DE50-MD dogs, (Figure 8). The LV pathology score and number of segments demonstrating fibrous or fibrofatty replacement of the myocardium both showed a highly significant linear relationship with the number of LGE positive segments (R^2^= 0.973, p<0.0001 and R^2^=0.939, p<0.0001) (figure 9). The number of segments detected in pathological sections tended to exceed those detectable by LGE by 1-2 segments. On post hoc cross-sectional comparison of images, LGE was able to detect fatty replacement with high accuracy, but occasionally failed to identify minimal (grade 1) fibrosis in the septum or apical regions, (Figure 11). Identification of small, linear bands of fibrosis in the septum was particularly challenging by LGE due to the potential for false interpretation of intramural, septal coronary vasculature in this region. Although Purkinje fibre vacuolation was present in DE50-MD dogs, this was not considered to be beyond that expected as a typical background lesion in healthy laboratory beagles [75]

**Figure 11:**
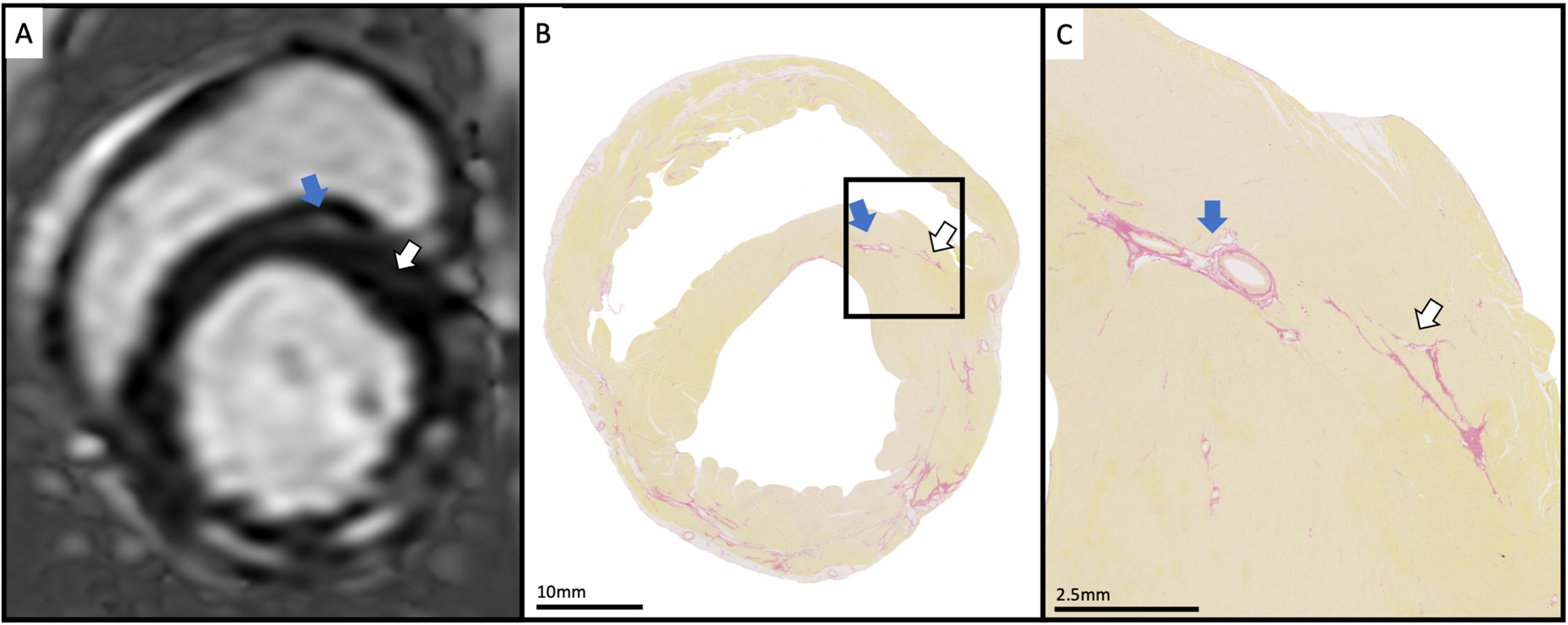
Example of cross-sectional imaging study showing late gadolinium enhancement (LGE) imaging (A), and corresponding picrosirius red histology images of the basal section of the heart of a LGE positive DESO-MD dog (Band C). In this dog, LGE was not identified prospectively in the septum. A single focus of grade 1 (minimal) fibrosis was identified in the anteroseptal segment (white arrow), adjacent to a septal intramural coronary artery (blue arrow). The septal vessel can be identified on the corresponding LGE image, but the presence of focal fibrosis identified on histopathology is much less clearly visible on the same image, (B and C).

## Discussion

Herein, we describe the progression of the preclinical cardiac phenotype in DE50-MD dogs aged 15-36 months of age. These data reveal that the left and right ventricular myocardium of DE50-MD dogs undergo progressive scarring and expansion of the ECM and that these changes are associated with systolic and diastolic functional decline, recapitulating the pathological progression described in DMD cardiomyopathy. Importantly, our study reveals the association between cardiac imaging, cTni and histopathological changes which hitherto, has been lacking in human DMD studies.

Whilst no DE50-MD dogs demonstrated overt systolic dysfunction, (as defined by a threshold LV EF of less than 55%), LV EF and average peak circumferential strain declined over the 12-to-15 months period from 15 months of age. Markers of RV function (TAPSE and RV S’) also declined over the same period. The age at which overt systolic dysfunction would be expected to occur in this model remains unclear but the changes documented are analogous to the rate of decline reported in the hitherto most comprehensively studied canine model of DMD: the Golden Retriever with Muscular Dystrophy (GRMD) dog model. Dogs from the GRMD dog colony in Texas[79] (described by Guo *et al*.) demonstrate a similar decline in average peak circumferential strain between 22 and 34 months of age but their LV EF did not significantly change over the same period. The dogs from that study, were followed to an older age (up to 77 months). The authors predicted a decline in EF to 54% at approximately 34 months of age and to below 40% (the cut-off specified to identify dilated cardiomyopathy in dogs)[80] at 43 months of age.[79] Systolic function was preserved in the DE50-MD dogs at 36 months, but direct comparison with the GRMD population is limited by the small number of dogs (n=4) remaining in our study at this age. Despite the decline in systolic function, cardiac dimensions and circulating NT-proBNP concentration remained lower in DE50-MD dogs compared to age matched WT dogs. These results mirrored those reported in young DE50-MD dogs and potential explanations have been reported elsewhere[63].

There was a high cumulative frequency of DE50-MD dogs demonstrating LGE over the study period. The overall prevalence and numbers of affected wall segments increased with age, but there was variation regarding age of onset and scar burden amongst individuals. Likewise, LGE positivity appears to be an age-related finding in DMD boys, [2, 4, 5, 11] with similar heterogeneity regarding age, being detectable from a very young age (<10-14 years of age) in some patients.[1, 4, 11] Late gadolinium enhancement corresponded to regional fibrofatty scarring of the myocardium in the majority of our cases. Rare, minor discrepancies likely related to insensitivity of LGE imaging to detect minimal fibrosis, false exclusion of LGE positive segments due to concern regarding misinterpretation of coronary vasculature and the fact that sections represented only an approximation of the imaging planes obtained during CMR imaging. In one dog, LGE accumulation corresponded to regions of active necrosis and oedema. The latter lesions were not encountered as commonly in the dogs studied here as in the GRMD dogs reported by Schneider *et al*.[78] In GRMD dogs, the appearance of myofibrillar necrosis precedes later chronic fibrofatty replacement in regions of myocyte loss.[78, 81] Similarly, myocardial necrosis was only identified in the youngest DE50-MD dog studied, which did not have evidence of fatty infiltration. Based on the combined results of our CMR and histopathological studies and the previous work ([78, 81]), we speculate that lesions progress from early regions of acute necrosis and inflammation to fibrotic replacement and then, more latterly, to fatty infiltration. Inclusion of a greater number of cardiac specimens encompassing a wider age range would be needed to test this hypothesis; however, if correct, it suggests that cardiac pathological changes mimic those detected in skeletal muscle [62].

In the DE50-MD dogs, LGE and fibrofatty scar distribution appeared in the subepicardial layer in the LV FW, as reported in early preclinical dystrophic cardiomyopathy.[4, 11, 82] Where present, LGE was seen in lateral LV segments, as in people with DMD,[12] but (predominately) fibrosis was also present in the septum of a subset of dogs. Early septal involvement is also described in GRMD dogs [79], but in DMD boys, septal segments tend to be spared from scarring until later in the disease course when there is a high fibrotic scar burden and further decline in LVEF.[4] As reported in GRMD dogs, [78, 81] DE50-MD dogs also had RV involvement histopathologically, that increased in severity alongside LV pathology. These findings coincide with their documented decline in RV function between 15 and 30 months of age. In people, assessment of the RV wall using CMR is challenging due to its comparatively thin lateral wall[83] and the presence of RV intramyocardial fat in healthy hearts.[84] The latter might also complicates the interpretation of our histopathological findings in the absence of age-matched controls. Early pathological descriptions of DMD cardiomyopathy report sparing of the RV from fibrosis,[36] but more recently, RV native T1 values were reportedly increased in DMD children, supporting the presence of diffuse fibrosis within the RV.[85] Further investigation in DE50-MD dogs could include parametric mapping studies of the RV in affected and control dogs and to further corroborate these findings with the results of histopathology.

Extracellular volume quantification had a strong association with the burden of LGE in the DE50-MD dogs studied and significantly exceeded that of WT dogs at all ages. The difference increased with age, suggesting an age-related progression of fibrosis and supporting the concept of diffuse ECM expansion prior to development of focal regional fibrosis. Measurement of ECV failed to demonstrate differences in myocardial composition in a longitudinal study of young GRMD dogs versus WT controls, (followed up to 12 months of age).[86] These results concur with pathological findings reported from the same colony, which demonstrated minimal cardiac fibrosis in GRMD dogs between 6 and 12 months.[78] Further investigation is therefore warranted at earlier ages to establish when differences in ECV can first be identified in the DE50-MD dog model. Global native myocardial T1 values of DMD boys with LGE exceed those of healthy controls in patients.[33, 34, 42] but there is conflicting information regarding the ability of T1 values to distinguish between LGE negative patients and controls.[34, 40, 42] The difference in native T1 times between DE50-MD and WT dogs did not become significant until 30 months of age, once values were corrected for confounding patient factors. Unfortunately, the variables with the greatest impact on native T1 were myocardial thickness, (which could be at least in part due to partial volume effects) and heart rate. These factors are particularly pertinent in this model due to increased heart rate and reduced heart size identified in DE50-MD dogs. We attempted to minimise the impact of these effects by careful measurement, avoiding the subendocardium and subepicardium and false inclusion of the blood pool or epicardial fat in native T1 measures, optimising breath holding protocols and imaging during the diastolic period to minimise cardiac movement. Further understanding regarding the effect of heart rate on the T1 measurement protocol could be gained through T1 mapping acquisition of imaging phantoms at variable heart rates. We did not attempt to quantify diffuse fibrosis in the interstitium of affected dogs, but comparison of parametric data from these dogs with the diffuse myocardial changes of the collected pathological specimens could help to further validate the use of these techniques.

The burden of fibrosis, (the number of LGE positive segments) and global ECV value, had a negative association with systolic and diastolic functional parameters in the DE50-MD dogs. Similarly, increasing percentage mass of LGE positive LV myocardium,[14, 82] increased LGE score[2] and transmurality[3] of LGE are associated with decline in EF and the number of LGE positive segments is an independent predictor of EF% [5] and mortality[4] in DMD patients. The relationship between LGE and circumferential strain parameters in DMD boys is less predictable. Circumferential strain deteriorates after development of LGE, even when EF is preserved[87, 88]but the decline appears to be regionally dependent. For example, deterioration of circumferential strain is more likely in the septum than in FW segments in the absence of LGE scarring.[12] In the DE50-MD dogs in this study, the number of LGE segments identified had a negative association with strain and systolic and early diastolic strain rate and the strength of association between LGE burden and peak circumferential strain was slightly higher than that with EF. Myocardial ECV is increased in DMD patents with EF<55%,[41] but its relationship with systolic function is not as clear as for LGE burden. Studies by Florian *et al*[89] and Soslow *et al*[40] both demonstrated that myocardial ECV had a significant negative correlation with EF in patients with (Duchenne and/or Becker) muscular dystrophy but these findings were not replicated in a later, larger study.[20]The association between functional parameters and ECV was weaker than that for LGE burden in the DE50-MD dogs. Regional ECV was only significantly associated with the early diastolic circumferential strain rate of associated segments. The association could be demonstrated, (albeit weakly) in the septum as well as FW segments, despite the lower frequency of scars in this region. There was a trend for a negative correlation between FW regional ECV and systolic parameters, (as has been shown in DMD boys[90]) but this association did not achieve statistical significance.

Measurement of native T1 times is appealing since contrast administration is not required and there is no delay for post-contrast imaging, shortening MRI acquisition times.[91] Despite the higher T1 values of DE50-MD dogs, average native T1 times were not associated with LV EF, as reported in DMD boys.[20, 33] T1 values were also unrelated to midventricular circumferential strain values, which is contrary to reports in people,[91] in whom T1 measurements correlate with CMR evaluation of circumferential strain. The evolution of change of native myocardial T1 values in DMD cardiomyopathy appears complex since older DMD patients, and those with advanced disease, (i.e. those with LGE and declining systolic function) can demonstrate unexpectedly low T1 values (<900ms).[20, 40] The presence of myocardial fatty replacement, (lipomatous metaplasia found in advanced disease) could provide a plausible explanation for these low values due to the very low native T1 values inherent to fat (250-300ms at 1.5T),[20] and complicate the interpretation of native myocardial T1 times in DMD cardiomyopathy.

In people, neither cTnI nor T1 times show a linear relationships with disease severity. In one study, the greatest cTnI plasma concentrations were measured in patients with only mild LGE and did not increase with patient age or with decline in ejection fraction.[92] Further, acute peaks in cTnI are recognised in DMD boys, sometimes coinciding with reported chest pain.[13, 93] This is supportive of episodes of recurrent myocardial injury and inflammation (myocarditis) in the dystrophin-deficient myocardium, which is important in the development of myocardial fibrofatty scarring. Likewise, three individual DE50-MD dogs showed transient severe elevations of cTnI that later plateaued to lower values (albeit still exceeding those of WT dogs). Even after exclusion of the outlier cTnI data, DE50-MD dogs had significantly higher cTnI that WT, and further, cTnI was higher in DE50- MD dogs demonstrating LGE compared to those without. Interestingly however, there was one outlier individual dog that did not develop LGE or show evidence of fibrofatty scarring or inflammation at post-mortem assessment. Neither cTnI or T1 values are sufficient as stand-alone measurements to replace post contrast imaging and reliably predict LGE status in DMD boys or DE50-MD dogs.[91, 92]

Measurement of T2 times should help to identify fatty tissue and oedema in DMD cardiomyopathy but there is a relative paucity of information in the literature regarding its measurement in this setting, compared to native T1 times and ECV. Measurement of T2 times did not discriminate between WT and DE50-MD dogs, which parallels findings from young DMD patients (<12 years of age).[54, 94] Unexpectedly, Soslow *et al* demonstrated that myocardial T2 times were, in fact, shortened in DMD boys compared to healthy controls.[40] High T2 values in some segments with LGE supported the presence of fatty infiltration or oedema in these individual patients, however. High T2 values were also obtained from some segments with LGE in individual DE50-MD dogs. These segments were those most frequently associated with fibrofatty infiltration in [76]DMD boys,[4, 11, 20, 36] GRMD dogs[78] and young DE50-MD dogs[63]. Although average midventricular T2 values failed to predict global functional derangements in the DE50-MD dogs, FW T2 times were associated with regional FW systolic dysfunction, as determined by composite peak circumferential strain values in these segments. The presence of regional fatty infiltration or oedema therefore appears to be associated with advancing disease when systolic function begins to decline. These findings are comparable to an early study by Wansapura *et al* of DMD boys.[94] In that study, the heterogeneity of T2 measurements significantly correlated with circumferential strain and ejection fraction. There was a strong association between T2 values and cTnI data in DE50-MD dogs, which further supports that cTnI release could be at least partly explained by myocardial inflammation, with resultant oedema and elevation in myocardial T2 times. Despite the overall similarity of T2 measurements between WT and DE50-MD dogs, the inclusion of T2 mapping alongside native T1 and ECV mapping provides additive information for myocardial characterisation and for monitoring myocardial changes in individual dogs.

All MRI markers of altered myocardial fibrosis (ECV, T1 and number of LGE positive segments) were significantly associated with the presence of fQRS on 12-lead echocardiographic assessment. The precise mechanism behind the presence of fQRS is not confirmed but is speculated to represent heterogeneity of electrical conduction within the ventricular myocardium, typically due to scarring or fibrosis.[95–97] In DE50-MD dogs, the number of leads demonstrating fQRS showed a highly significant negative association with markers of global and midventricular circumferential systolic, (EF, circumferential strain and systolic strain rate) and diastolic, (early diastolic circumferential strain rate) function. Similar findings are reported in children with DMD[98] who develop fQRS from a young age.[72] In a small study of 37 DMD boys aged 6-28 years, (15y±5.5) the presence of fQRS was associated with lower EF, increased Tei index (a marker of combined global systolic and diastolic function) and increased burden of ventricular arrhythmia. Further, the number of leads with fQRS showed a negative correlation with longitudinal myocardial systolic and diastolic velocities at the level of the mitral anulus, EF and VPC burden and a positive correlation with global longitudinal strain, VPC burden and Tei index. Unlike DE50-MD dogs, in DMD boys enrolled in the study by Cho *et al* there was no association between fQRS and LGE burden. These contrary findings are likely explained by the different methods used to quantify LGE (% area in that study rather than number of segments with LGE). The appearance of fQRS complexes in the inferior, (leads II, III and aVF) and lateral leads, (leads I and aVL)[99] parallels the high frequency of LGE accumulation in inferior and lateral wall segments in the DE50-MD dogs and is to our knowledge, previously unreported in an animal model of DMD; fQRS assessment in in DE50-MD dogs appears to be a simple assessment that can be performed in conscious DE50-MD dogs, corroborating the results from CMR markers of fibrosis severity in this model.

## Study limitations

The main limitation of this study is the lack of conscious functional echocardiographic data from WT dogs due to their poor tolerance for lateral recumbency, restricting meaningful longitudinal comparisons of functional parameters (in particular those relating to STE and TDI techniques and of diastolic function parameters). Sedation was considered, as has been described during cardiac assessments in other DMD animal models[100] but this was avoided due to the concern regarding the potential for unmeasurable differential effects of sedation between WT and DE50-MD dogs. Several DE50-MD dogs required euthanasia prior to the planned study end point, (36 months), which complicated interpretation of some non-statistical analyses (for example, longitudinal evaluation of functional parameters). Further, the small number of DE50-MD dogs studied precluded further subgroup analysis between those with and without LGE; for example, exploration of differences in parametric mapping and LV function between dogs with and without LGE, and this could be an area of future study. Although there was excellent corroboration of LGE imaging and severity of fibrofatty lesions within the LV, the interval of 66 and 83 days respectively between the final CMR and date of euthanasia in two dogs could have affected our findings. Finally, our histopathological analysis was not designed to morphometrically quantify interstitial myocardial changes and so we did not attempt to compare results of parametric mapping studies with histopathological data. This also could be considered in future studies.

## Conclusions

The data presented in this study demonstrate that there is continued progression of the cardiac phenotype in DE50- MD dogs from 15 months of age that resembles that of DMD children. The rate of progression in this model appears to be similar but overall, is slightly more benign than that of the GRMD colony in Texas. Direct comparison between models is challenging however as, to date, no DE50-MD dog has been studied to the late ages of the dogs studied from the Texas colony. Nevertheless, the results of conscious echocardiography demonstrated functional decline in LV EF and RV functional data, in addition to circumferential strain parameters in DE50-MD dogs that was associated with blood borne (cTnI) and CMR markers of diffuse and focal myocardial pathology. The pattern and distribution of myocardial scarring determined using histopathology and cross sectional LGE imaging resembled that of GRMD dogs and DMD boys. The cumulative frequency of LGE was high in the DE50-MD dogs by 36 months of age, but where present, it was principally limited to the subepicardial location, with no dogs showing evidence of transmurality in the LV. Septal involvement appears to be relatively common feature of canine dystrophic cardiomyopathy, which differs from findings in DMD children. Likewise, RV pathology was also seen alongside LV pathology in DE50-MD dogs.

Parametric mapping techniques were feasible in DE50-MD and WT dogs and exposed diffuse myocardial pathological changes that were most pronounced in DE50-MD dogs of 30 months and older. The combined results of T1, ECV and T2 mapping studies were supportive of ECM expansion due to fibrosis, with or without regional myocardial oedema and /or fatty infiltration in individual dogs. This was corroborated by the results of cTnI and histopathological analysis. The presence of fQRS complexes in DE50-MD dogs was associated with MRI markers of fibrosis and deterioration in LV function and provide a potentially simple, non-invasive method with which to monitor disease progression in this model. The results from this study, in combination with our previous work [63] provide a comprehensive description of the preclinical phenotypic progression of the DE50-MD model and are useful to establish pr otocols for myocardial characterisation of the model for use in ongoing clinical trials. Our work reveals the associations between the characteristic myocardial structural and functional decline and contemporaneous histopathology which has hitherto, been challenging in human DMD patients, emphasizing the utility of this animal model.

## Acknowledgements

The authors thank colleagues at the Royal Veterinary College for their contributions and especially Dr Yu-Mei Chang for advice on statistical analysis and staff within the Biological Services Unit for excellent and compassionate care of the animals. This research was funded by the Wellcome Trust (101550/Z/13/Z) and by Muscular Dystrophy UK (RA3/3077/2).

