## Supplementary information for "Progression of the cardiac phenotype of the DE50-MD dog model of Duchenne Muscular Dystrophy, corroborating results of cardiac magnetic resonance imaging with pathology up to 36 months of age"

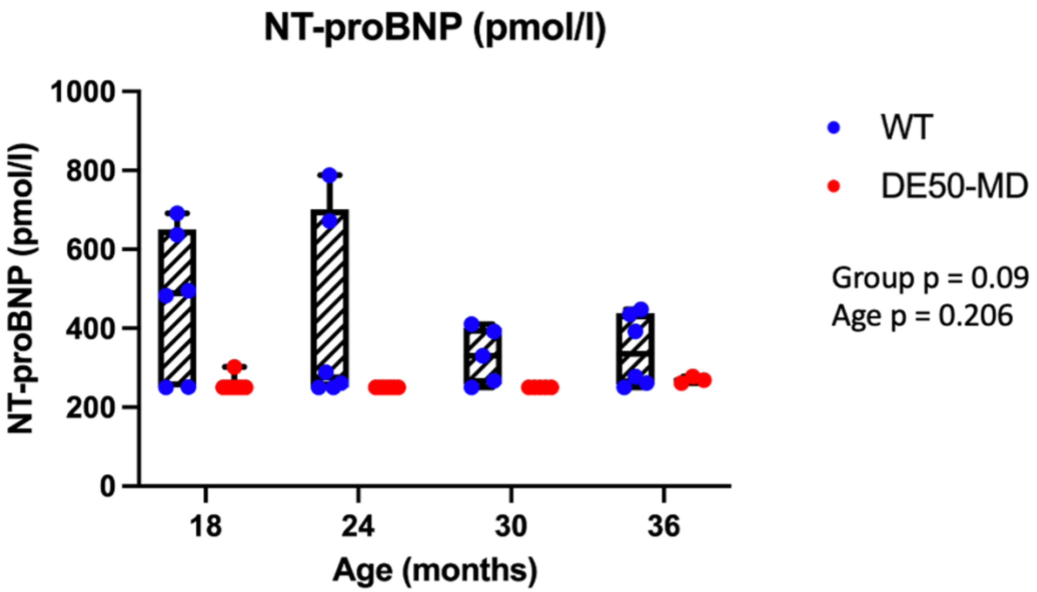


***Supplementary figure 1: Plasma NT-proBNP concentration in DE50-MD and WT control dogs. NTproBNP concentration (pmol/l) in the plasma of DE50-MD (red) and WT dogs (blue) DE50-MD: total n=8 dogs, n=1-4 per age group; WT: total n=6 dogs, N=3-4 per age group****. The lower limit of detection of the assay is 250pmol/l. Each dot represents an individual dog. Boxes extend from the 25th to 75th percentile, with a line within the box at the median value. Each point represents an individual dog, and whiskers show the minimum and maximum results for that age-group. DE50-MD dogs had lower circulating NT-proBNP, with no significant effect of age.*

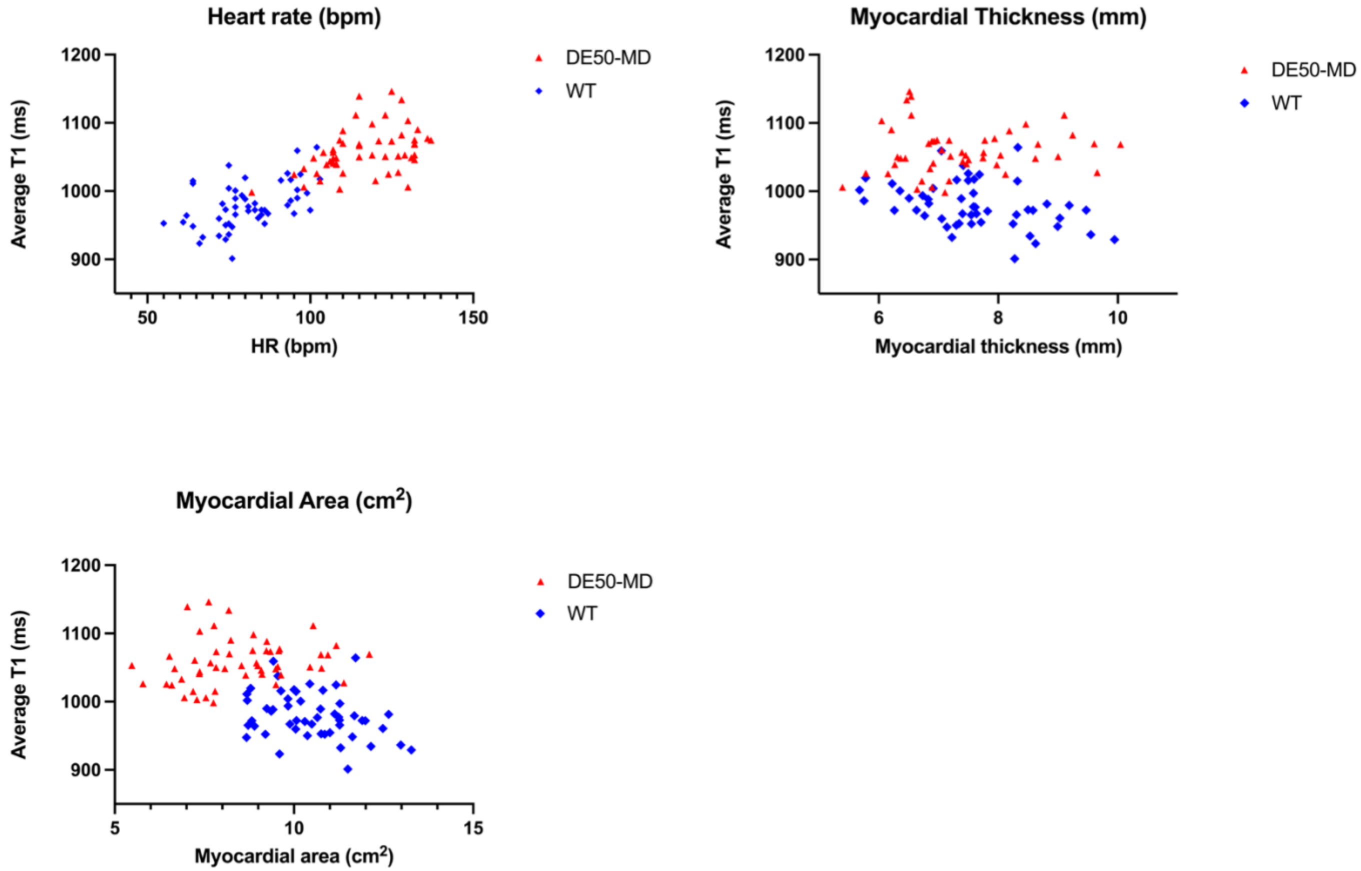


***Supplementary figure 2: Relationship between i) heart rate (HR), ii) myocardial thickness and iii) area with average native T1 times in the base and midventricular slices for DE50-MD and wild type control (WT) dogs.*** *A linear association was demonstrated between average T1 and HR for both WT and DE50-MD dogs (r=0.434 and 0.501, p=0.0013 and p<0.0001 respectively). Significant but very weak linear associations between average T1 values and myocardial area and wall thickness were only identified in WT dogs (r=-0.304 and r=-0.370; p=0.030 and 0.008 respectively).*

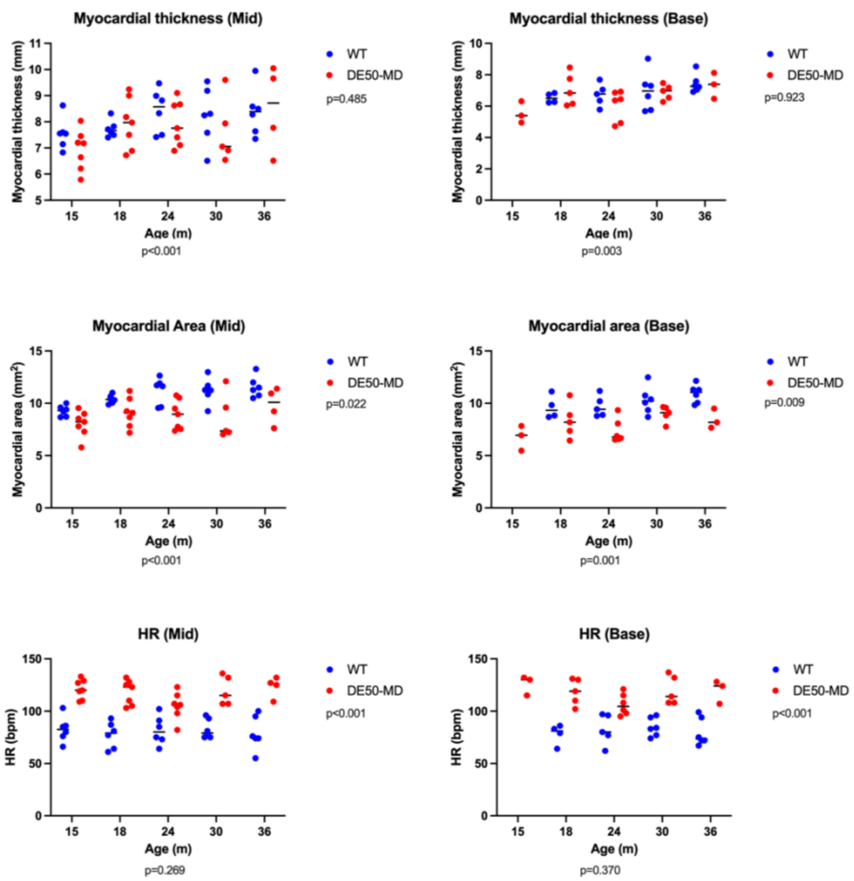


***Supplementary figure 3: Myocardial thickness and area, and heart rate recorded during assessment of average native T1 times in the base and midventricular slices for DE50-MD (total n=8 dogs, n=3-7 per age group) and wild type (WT, total n=6, n=4-6 per age group) dogs.*** *On evaluation of both the base and midventricular slices, myocardial area was reduced (p=0.009 and p=0.022 respectively) and heart rate was increased (p<0.001 for both) in DE50-MD dogs compared WT dogs. Myocardial thickness was similar between genotypes for both ventricular levels (p=923 and p=0.475 respectively).*

**Transthoracic Echocardiography protocols:**

| Dimension | View | Description of technique | Timing in cardiac cycle | Method of normalisation |
|---|---|---|---|---|
| **LV volume:** | RPx4ch | Estimated using Simpson's method of discs by tracing the endocardial border of the LV at end diastole and systole. | LV end diastolic volume (LVEDV): first frame after mitral valve closure.<br>LV end systolic volume (LVESV): last frame prior to mitral valve opening | |
| **LV internal diameter** | RPxSA | RPxSA views at the level of the chordae tendinae. M Mode recording, with the cursor bisecting the LV. Measured using a leading edge to leading edge technique. | LV end diastolic internal diameter (LVIDd), septal (IVS) and free wall (FW) diameters: at the onset of the QRS complex.<br>LV end systolic internal diameter (LVIDs): at the nadir of septal excursion. | According to body weight using Esser et al allometric scaling method:<br>LVIDDN: LVIDD/BW $^{0.322}$<br>LVIDSN:LVIDS/BW$^{0.346}$ |
| **Tricuspid Annular Systolic plane excursion (TAPSE)** | Left apical | M-mode recordings with the cursor as parallel as possible to the majority of the RV free wall. Measurement of the maximal longitudinal displacement of the lateral tricuspid valve annulus toward the RV apex | Peak systole. | According to bodyweight: TAPSE/ BW$^{0.297}$<br>2. According to aortic diameter (AV diam)<br>LVIDd:Ao and LVIDs:Ao |

**Calculations:**

Ejection Fraction (EF): ((LVEDV-LVESV)/LVEDV) x 100

Fractional Shortening (FS): ((LVIDd-LVIDs)/LVIDd) x 100

**12 lead Electrocardiography protocol:**

Studies were performed directly before and after conscious echocardiographic assessments using the same electrocardiography machine (Nihon Kohden Cardiofax V 1550K).

Dogs were gently restrained in right lateral recumbency with thoracic limbs held perpendicular to the long axis of the body and pelvic limbs held in a semiflexed position. The leads were recorded according to standardised protocols for dogs.

Bipolar limb leads:

I: Right thoracic limb (-) to left thoracic limb (+)

II: Right thoracic limb (-) to left pelvic limb (+)

III: left thoracic limb (-) to left pelvic limb (+)

Augmented unipolar lead

aVR: Averaged signal from left thoracic limb and left pelvic limb (-) to right thoracic limb (+)

aVL: Averaged signal from right thoracic limb and left pelvic limb (-) to left thoracic limb (+)

aVF: Averaged signal from right and left thoracic limb (-) to left pelvic limb (+)

Precordial leads were placed as follows:[765]

V1: adjacent to the sternum in the 1st intercostal space at the level of the costochondral junction

Electrodes for the precordial leads V2 through V6 were positioned along the 6th intercostal space as follows:

V2: adjacent to the sternum

V3: midway between V2 and V4,

V4: at the costochondral junction

V5 and V6: dorsal to V4 at a distance equal to that between V3 and V4

The electrocardiogram was recorded at a 50mm/second paper speed for a least 1 minute.

**Cardiac Magnetic resonance Imaging Protocol:**

**Standard imagine sequences:**

Two- and 4-chamber VLA cine images were prescribed from survey gradient echo transverse, sagittal and dorsal standard localising scans (scouts). Cine image acquisition was performed using a gradient echo pulse sequence with balanced steady-state free precession (bright-blood imaging). A series of short axis images of the LV was obtained using the 2- and 4-chamber VLA sequences. Care was taken to ensure that the image plane was parallel to the plane of the mitral annulus and that acquired images encompassed the entire LV and LA from the apex of the LV to the dorsal aspect of the LA. Contiguous 5mm slices were acquired, with no interslice gap and 30 frames per cardiac cycle.

Cardiac magnetic resonance images were viewed using open source Osirix software (http:// www.osirix-viewer.com). Left ventricular mass, end-systolic and end-diastolic volumes were obtained using the disc summation method. Manual contours were drawn along the endocardial and epicardial borders from the end systolic and end-diastolic frames of the short axis stack. The end-diastolic image was defined as the frame with maximum dilatation and the end-systolic image as the frame with maximum contraction during the recorded cine loops. Papillary muscles were excluded but LV trabeculae were included in the ventricular lumen volume. The most basal slice included was the slice where the endocardial contours were interrupted by the LV outflow (even at end diastole) but greater than 50% of the ventricular lumen circumference was surrounded by a wall thickness consistent with ventricular myocardium. The LV mass was calculated both in systole and diastole by subtracting the endocardial volume from the epicardial volume and multiplying the result by the myocardial mass density (1.05g/ml).[644, 645] Where possible, contouring of the RV lumen was repeated in both the transverse and the short axis stack imaging planes. The equivalent frames to those selected for LV volumetric measurements were used for end-systolic and end-diastolic volume calculation, with the tricuspid and pulmonary valves used as the limits for right ventricular volume. Only endocardial contouring was performed, excluding the papillary muscles from RV blood volume calculation for consistency.

Left ventricular and RV stroke volumes were calculated by subtracting calculated end systolic ventricular volumes from end diastolic ventricular volumes. Measurements for RV volume were accepted if calculated RV stroke volumes from transverse and short axis imaging planes were within 10% agreement. Likewise, measurements for LV volumes were accepted if values for systolic and diastolic mass were within 10% agreement. Average values were reported for LV mass and for RV volumes. Ventricular volumes and LV mass were normalised to patient size by indexing calculated values to body surface area (BSA) to yield the following variables: indexed LV end diastolic volume (LVEDVI) and indexed stroke volume (LVESVI).

| | 15 | | 18 | | 24 | | 30 | | 36 | |
|---|---|---|---|---|---|---|---|---|---|---|
| ECV | DE50-MD | WT | DE50-MD | WT | DE50-MD | WT | DE50-MD | WT | DE50-MD | WT |
| BASE | | | | | | | | | | |
| inferior | 21.5 (17.2-22.2) | | 22.6 (20.0-24.5) | 20.3 (18.9-23.0) | 21.0 (19.7-25.7) | 18.7 (17.3-20.2) | 21.8 (19.0-24.0) | 17.6 (16.7-18.9) | 22.5 (19.9-23.1) | 18.2 (16.7-21.0) |
| inferolateral | 19.8 (16.7-20.2) | | 21.0 (18.8-22.8) | 18.7 (17.3-21.1) | 19.2 (18.2-28.8) | 17.6 (16.1-18.0) | 21.0 (18.4-25.9) | 16.7 (15.4-17.0) | 21.1 (19.6-26.9) | 17.4 (15.4-22.0) |
| anterolateral | 18.0 (17.1-18.6) | | 20.5 (18.9-24.3) | 17.9 (16.0-19.1) | 20.2 (17.9-26.6) | 17.0 (16.4-18.2) | 19.3 (17.4-32.0) | 16.1 (15.2-16.4) | 26.1 (19.5-28.5) | 16.4 (14.5-17.8) |
| anterior | 17.9 (16.0-19.6) | | 21.5 (19.2-25.0) | 19.1 (16.8-20.3) | 21.3 (19.8-24.7) | 18.5 (17.7-19.9) | 20.4 (17.8-27.8) | 17.1 (15.9-17.8) | 22.1 (19.2-26.6) | 17.5 (14.8-18.4) |
| anteroseptal | 22.9 (19.7-25.3) | | 25.4 (21.6-28.1) | 21.4 (19.5-27.4) | 27.6 (25.0-29.3) | 23.2 (20.2-25.9) | 24.1 (20.4-26.8) | 21.3 (18.6-22.9) | 26.3 (23.8-26.6) | 21.6 (18.4-24.7) |
| inferoseptal | 20.5 (19.4-22.2) | | 23.2 (22.3-29.4) | 20.2 (19.4-20.9) | 23.8 (21.2-25.9) | 19.6 (18.4-20.2) | 21.5 (21.0-25.9) | 18.7 (16.6-20.5) | 24.7 (21.7-24.7) | 18.7 (16.9-20.6) |
| MID | | | | | | | | | | |
| inferior | 20.9 (19.8-23.0) | 20.7 (18.6-22.7) | 22.0 (19.6-25.6) | 19.6 (17.4-20.6) | 22.3 (19.5-25.6) | 18.3 (17.0-19.6) | 22.4 (18.3-24.1) | 18.0 (16.7-18.8) | 21.8 (19.8-28.4) | 17.8 (16.6-20.8) |
| inferolateral | 19.3 (18.0-26.7) | 18.6 (17.4-19.8) | 21.8 (17.8-30.5) | 18.1 (15.9-19.4) | 20.3 (18.1-25.2) | 17.0 (15.4-17.8) | 22.3 (17.1-23.3) | 16.7 (15.2-17.8) | 21.2 (19.5-28.0) | 16.4 (14.4-17.8) |
| anterolateral | 19.8 (18.7-26.1) | 18.8 (17.7-19.9) | 21.8 (19.1-28.5) | 19.1 (15.8-19.7) | 20.4 (18.1-28.5) | 17.2 (16.8-19.6) | 22.6 (17.6-23.4) | 16.5 (15.6-17.9) | 20.3 (19.3-30.0) | 17.0 (15.0-17.5) |
| anterior | 20.2 (17.7-23.5) | 18.9 (18.5-19.3) | 22.3 (20.1-25.0) | 20.1 (16.9-21.1) | 21.7 (18.9-24.9) | 17.5 (16.9-19.6) | 21.2 (18.7-24.3) | 17.6 (16.6-18.9) | 20.6 (19.7-27.4) | 17.3 (15.4-19.0) |
| anteroseptal | 22.9 (19.0-23.7) | 20.4 (19.2-21.5) | 21.8 (19.4-23.0) | 20.1 (17.8-21.7) | 21.8 (18.7-29.4) | 18.1 (17.4-19.3) | 22.9 (18.5-23.5) | 17.9 (16.6-18.7) | 21.9 (19.2-27.3) | 18.3 (16.4-19.7) |
| inferoseptal | 21.8 (19.4-25.3) | 20.6 (19.1-22.0) | 23.3 (20.0-24.0) | 20.2 (19.2-21.2) | 22.9 (17.9-29.0) | 17.9 (17.5-18.8) | 23.7 (19.2-25.2) | 18.7 (17.3-18.9) | 23.9 (21.2-28.2) | 19.1 (15.7-20.1) |
| APEX | | | | | | | | | | |
| inferior | 22.3 (18.9-22.7) | | 23.5 (20.0-24.6) | 19.6 (17.9-20.9) | 22.2 (20.5-25.6) | 18.8 (18.2-19.5) | 20.1 (19.7-23.1) | 18.0 (16.2-20.0) | 21.4 (20.0-25.1) | 18.9 (15.8-20.0) |
| inferolateral | 21.1 (18.5-22.3) | | 22.3 (19.4-23.8) | 19.2 (16.5-21.4) | 20.5 (18.4-24.0) | 18.3 (16.8-18.6) | 19.5 (17.6-22.3) | 17.6 (15.0-19.1) | 19.9 (17.7-22.3) | 18.0 (15.5-19.1) |
| anterolateral | 20.7 (19.8-24.0) | | 24.5 (21.6-26.2) | 19.3 (18.2-20.5) | 22.1 (20.6-25.4) | 18.4 (17.7-19.7) | 20.8 (18.9-23.5) | 18.8 (17.2-19.2) | 20.4 (18.9-23.5) | 18.3 (17.1-19.9) |
| anterior | 21.4 (19.5-23.4) | | 23.6 (22.4-26.8) | 20.8 (18.7-21.4) | 22.8 (20.3-24.7) | 19.8 (19.1-20.4) | 20.9 (18.5-24.6) | 18.2 (16.6-19.5) | 21.6 (19.3-23.6) | 18.6 (15.9-19.1) |
| anteroseptal | 21.0 (19.7-22.0) | | 23.8 (20.8-25.6) | 20.7 (19.7-21.3) | 23.1 (21.6-25.3) | 19.8 (18.0-21.1) | 23.5 (19.8-25.1) | 18.8 (17.9-20.4) | 22.6 (19.2-24.3) | 19.0 (16.9-20.9) |
| inferoseptal | 22.7 (20.9-26.0) | | 23.1 (21.0-27.1) | 20.7 (18.5-21.6) | 23.4 (21.3-27.1) | 19.0 (18.2-20.1) | 23.7 (18.9-26.0) | 19.2 (14.8-20.3) | 22.5 (20.4-23.3) | 18.8 (15.7-21.2) |
| T2 | | | | | | | | | | |
| inferior | 45.1 (43.0-49.3) | 44.3 (43.3-46.5) | 45.1 (44.6-46.5) | 45.7 (42.8-46.5) | 46.1 (44.6-54.9) | 45.5 (44.5-48.5) | 45.3 (43.4-50.1) | 45.4 (42.3-47.1) | 49.1 (45.8-52.2) | 45.2 (40.1-53.5) |
| inferolateral | 45.2 (43.1-48.8) | 44.5 (42.0-47.6) | 46.4 (42.7-53.2) | 42.9 (42.5-45.1) | 45.1 (42.7-51.5) | 45.1 (44.4-46.5) | 46.0 (42.0-47.9) | 45.6 (41.9-46.3) | 46.1 (43.7-48.6) | 43.2 (40.4-45.4) |
| anterolateral | 44.8 (40.0-47.2) | 44.0 (42.7-46.6) | 42.2 (41.2-46.7) | 43.2 (41.2-45.1) | 42.9 (41.5-48.0) | 44.7 (43.2-47.0) | 44.8 (40.9-45.3) | 42.7 (41.8-45.6) | 45.0 (43.3-47.1) | 42.9 (38.6-46.8) |
| anterior | 45.4 (41.9-48.3) | 45.4 (43.3-47.2) | 45.1 (39.9-47.9) | 43.3 (42.9-49.4) | 44.9 (41.5-48.4) | 45.6 (42.7-48.3) | 42.5 (40.8-46.8) | 45.9 (43.5-48.1) | 45.2 (43.7-47.0) | 44.7 (40.7-49.3) |
| anteroseptal | 47.3 (42.4-52.9) | 45.3 (43.8-47.2) | 47.7 (42.5-48.2) | 44.6 (42.1-49.4) | 45.1 (44.6-47.4) | 45.8 (44.9-56.6) | 45.5 (42.6-46.8) | 43.8 (41.8-46.8) | 47.0 (44.8-49.2) | 44.5 (39.5-46.0) |
| inferoseptal | 45.4 (42.3-49.1) | 45.2 (44.2-46.1) | 45.5 (39.9-48.6) | 43.7 (43.2-51.8) | 46.0 (44.2-53.7) | 47.8 (43.8-49.9) | 45.0 (42.8-46.9) | 47.5 (43.1-56.7) | 45.1 (43.4-47.0) | 44.4 (40.2-51.2) |
| Native T1 | | | | | | | | | | |
| BASE | | | | | | | | | | |
| inferior | 1065.8 (1014.5-1125.5) | | 1120.9 (1005.5-1226.3) | 995.6 (918.2-1011.5) | 1036.8 (923.6-1088.1) | 1005.9 (934.4-1104.7) | 1060.2 (1025.6-1188.0) | 970.5 (950.6-978.9) | 1033.7 (1004.7-1048.2) | 965.65 (913.4-1017.8) |
| inferolateral | 1047.7 (1012.8-1075.0) | | 1065.9 (1029.6-1122.3) | 988.9 (948.8-1007.6) | 1031.4 (922.4-1054.1) | 996.8 (938.1-1022.1) | 1070.2 (1048.1-1125.3) | 969.7 (949.9-986.1) | 1093.8 (1078.7-1218.3) | 965.9 (908.0-1014.0) |
| anterolateral | 1023.9 (1019.3-1032.2) | | 1041.0 (1001.9-1068.0) | 977.6 (970.4-1037.0) | 1027.6 (989.3-1108.6) | 986.4 (955.3-1031.6) | 1023.8 (1015.2-1055.5) | 966.7 (903.5-980.1) | 1030.8 (1023.8-1164.1) | 961.7 (930.3-1006.4) |
| anterior | 1005.9 (982.0-1071.4) | | 1055.8 (1030.3-1086.8) | 999.1 (978.7-1013.0) | 1089.7 (1032.9-1168.6) | 1054.1 (1010.9-1074.9) | 1068.6 (1020.1-1138.8) | 981.7 (948.9-1040.7) | 1066.0 (1004.7-1151.7) | 990.8 (921.5-1014.8) |
| anteroseptal | 1042.1 (1008.4-1073.1) | | 1082.1 (1005.6-1107.3) | 998.5 (980.5-1028.0) | 1084.6 (1054.2-1103.7) | 1047.9 (997.2-1108.3) | 1049.5 (1028.0-1146.9) | 988.5 (957.0-1050.7) | 1060.7 (1009.7-1182.6) | 1004.9 (958.7-1033.9) |
| inferoseptal | 1015.7 (997.8-1041.7) | | 1061.6 (1027.6-1111.4) | 985.0 (965.8-1004.0) | 1042.3 (1000.4-1070) | 1002.3 (939.2-1014.0) | 1054.3 (1053.4-1063.2) | 974.5 (946.5-1026.2) | 1041.7 (999.0-1082.7) | 972.7 (903.9-1045.3) |
| MID | | | | | | | | | | |
| inferior | 1046.0 (1001.1-1095.7) | 950.9 (905.2-1041.8) | 1054.5 (1017.6-1096.6) | 960.9 (950.3-1031.3) | 1057.2 (1014.6-1104.1) | 985.9 (937.8-1099.8) | 1079.8 (1051.9-1166.1) | 954.5 (926.9-986.9) | 1071.5 (1056.6-1112.1) | 944.3 (877.8-970.2) |
| inferolateral | 1035.3 (997.2-1137.4) | 955.5 (935.0-1012.1) | 1076.0 (1002.0-1168.4) | 966.5 (948.7-1025.9) | 1017.9 (999.2-1133.6) | 1001.2 (951.0-1053.5) | 1067.8 (1041.4-1227.5) | 957.6 (937.4-998.8) | 1076.7 (1061.3-1376.7) | 934.7 (900.0-966.4) |
| anterolateral | 1038.4 (996.4-1061.0) | 966.6 (926.3-998.1) | 1049.0 (1019.4-1144.7) | 975.0 (944.2-1066.3) | 1023.1 (970.3-1193.0) | 993.8 (961.1-1109.5) | 1068.4 (1042.1-1072.2) | 962.9 (936.5-997.5) | 1122.4 (993.2-1149.2) | 942.9 (916.5-988.7) |
| anterior | 1028.8 (996.9-1077.2) | 981.6 (922.6-1013.4) | 1069.9 (1040.5-1096.1) | 993.6 (947.3-1044.5) | 1060.2 (1010.7-1131.5) | 1013.0 (946.3-1076.3) | 1075.7 (1046.4-1154.8) | 990.8 (948.9-1007.1) | 1084.5 (1016.4-1105.4) | 960.4 (910.5-997.0) |
| anteroseptal | 1045.1 (987.6-1078.1) | 967.0 (916.6-1033.4) | 1034.3 (993.7-1063.1) | 974.8 (938.4-1018.8) | 1049.1 (983.2-1079.8) | 1010.7 (939.2-1072.4) | 1052.1 (1023.6-1121.8) | 966.4 (928.8-1005.0) | 1021.5 (997.6-1053.3) | 960.1 (878.9-1059.5) |
| inferoseptal | 1060.0 (989.4-1107.1) | 963.7 (907.2-1013.0) | 1062.1 (992.4-1086.1) | 990.3 (956.7-1027.7) | 1072.8 (969.1-1084.6) | 991.3 (935.5-1048.2) | 1050.5 (1011.2-1107.5) | 972.7 (938.4-994.0) | 1055.6 (1017.7-1097.8) | 964.1 (902.8-1043.9) |
| APEX | | | | | | | | | | |
| inferior | 1105.5 (1039.6-1128.1) | | 1092.4 (1039.4-1200.1) | 1008.5 (922.3-1093.3) | 1121.2 (998.9-1239.8) | 1058.1 (997.8-1122.5) | 1067.3 (1061.7-1122.7) | 994.5 (949.8-1035.1) | 1085.0 (1022.8-1122.2) | 955.0 (945.7-988.8) |
| inferolateral | 1085.7 (1047.6-1145.2) | | 1148.9 (1011.4-1209.4) | 1044.9 (1017.8-1135.1) | 1068.4 (1034.9-1134.0) | 1001.5 (962.8-1238.1) | 1063.4 (1053.7-1271.3) | 966.6 (942.3-1084.6) | 1060.8 (1039.3-1089.7) | 970.5 (909.8-1078.5) |
| anterolateral | 1078.3 (1061.1-1353.6) | | 1175.8 (1128.8-1217.0) | 1036.6 (950.8-1133.3) | 1171.8 (1085.9-1320.4) | 1056.1 (955.9-1105.3) | 1106.7 (1070.6-1188.5) | 1014.2 (950.1-1058.9) | 1219.4 (1050.2-1313.8) | 1001.4 (916.3-1123.8) |
| anterior | 1112.7 (1030.6-1370.4) | | 1245.9 (1206.0-1439.3) | 1027.2 (1016.1-1118.7) | 1231.7 (1107.1-1325.1) | 1052.4 (987.3-1104.2) | 1121.3 (1004.0-1181.3) | 1013.6 (933.0-1079.0) | 1198.2 (1020.1-1288.4) | 1004.2 (930.2-1109.5) |
| anteroseptal | 1050.0 (1008.6-1089.4) | | 1068.7 (1042.7-1120.6) | 1040.7 (970.0-1092.0) | 1082.9 (1030.6-1125.8) | 1026.5 (988.5-1103.3) | 1095.4 (1052.1-1155.4) | 1003.7 (956.0-1124.9) | 1054.2 (1040.5-1133.0) | 999.3 (959.7-1055.4) |
| inferoseptal | 1105.7 (1078.2-1146.7) | | 1072.4 (1015.5-1267.0) | 1009.5 (947.0-1024.9) | 1088.8 (1033.3-1141.5) | 1011.8 (965.3-1028.3) | 1052.6 (1011.2-1151.8) | 987.5 (921.3-1026.5) | 1047.6 (987.7-1125.1) | 977.1 (937.9-1020.4) |

Supplementary table 1: Results of parametric mapping studies, showing segmental median (range) T1 times, T2 times and extracellular volume (ECV) in Wild type (WT) and DE50-MD dogs
